# Kinematic analysis of cell lineage reveals coherent and robust mechanical deformation patterns in zebrafish gastrulation

**DOI:** 10.1101/054353

**Authors:** David Pastor-Escuredo, Benoît Lombardot, Thierry Savy, Adeline Boyreau, Jose M. Goicolea, Andrés Santos, Paul Bourgine, Juan C. del Álamo, Nadine Peyriéras, María J. Ledesma-Carbayo

## Abstract

Digital cell lineages reconstructed from 3D+time imaging data provide unique information to unveil mechanical cues and their role in morphogenetic processes. Our methodology based on a kinematic analysis of cell lineage data reveals deformation patterns and quantitative morphogenetic landmarks for a new type of developmental table. The characteristic spatial and temporal length scales of mechanical deformation patterns derived from a continuous approximation of cell displacements indicate a compressible fluid-like behavior of zebrafish gastrulating tissues. The instantaneous deformation rate at the mesoscopic level of the cell’s neighborhood is spatially and temporally heterogeneous. The robustness of mechanical patterns results from their cumulative history along cell trajectories. Unsupervised classification of mechanical descriptor profiles was used to assess the homogeneity of biomechanical cues in cell populations. Further clustering of cell trajectories according to their cumulative mesoscopic biomechanical history during gastrulation revealed ordered and coherent spatiotemporal patterns comparable to that of the embryonic fate map.

## Introduction

Morphogenetic processes shaping animal embryos depend on the mechanical properties of cells and tissues and the generation and transduction of forces^1,2^. Mechanical stimuli affect all levels of biological organization including gene expression and signaling activities underlying biological processes such as cell proliferation and apoptosis^3^. Cell stiffness, contractility, protrusive activity and adhesive properties are the basis for these cell and tissue mechanical properties. The robustness of embryonic development depends on the multiscale coordination of biochemical and biomechanical processes in space and time^4–6^. Recent advances in live microscopy imaging supporting the digital reconstruction of cell lineages and cell shapes^7–11^ opened the way to a quantitative and formal approach of cell dynamics and tissue deformation^12,13^. The quantitative analysis of cell displacements and shape changes from time-lapse imaging data has been used to characterize, in 2 dimensions (2D), tissue deformation rate patterns and their perturbation in mutants^3,13–15^. Modeling tissue properties and cell contacts in 2D^16,17^ or designing bulk constitutive equations^19^ has also been successfully used to evaluate stresses. However, the current challenge^4–6^ is to tackle processes in 3D over time and correlate biomechanical patterns with other spatiotemporal features including gene expression dynamics and cell fate determination^20^. Owing to its amenability to live in vivo imaging^7,21,22^ and fast embryonic development, the zebrafish is a valuable model organism to investigate the spatial and temporal dynamics of tissue deformation underlying vertebrate gastrulation^23,24^.

We developed a methodology to assess in 3D the instantaneous and cumulative tissue deformation based on digital cell lineage trees reconstructed from partial 3D+time imaging of gastrulating zebrafish embryos^25^. Descriptors were computed to quantify tissue compression or expansion, rotation and distortion at the mesoscopic scale of the cell’s neighborhood. The instantaneous deformation rates provided biomechanical landmarks to identify gastrulation phases. We observed that the spatial and temporal heterogeneity of these landmarks was compensated along cell trajectories through a cumulative process. The computational categorization of cumulative mechanical descriptors led to the definition of a limited number of Canonical Lagrangian Biomechanical Profiles (CLBPs). The biomechanical signature of each cell derived from CLBPs was used to build a Lagrangian Biomechanical Map (LBMap) revealing the order and coherence of mechanical cues and their correlation with cell fate.

## Results

### Instantaneous deformation descriptors provide landmarks for the progression of zebrafish gastrulation

We developed a computational framework to quantify strains at the mesoscopic scale of the cell and its neighborhood in digital cell lineage trees reconstructed from 3D+time imaging data of developing embryos^22^. Our methodology was applied to five zebrafish embryos (z1-z5) imaged throughout gastrulation from the animal pole, to observe the development of their presumptive brain with a 2.5 min temporal resolution, and a voxel size of 1.4 μm^3^ (Supplementary Fig. 11, Supplementary Movie 1, Supplementary Table 1). Mechanical patterns and their evolution in space and time were revealed through tensorial analysis of the continuous approximation of the cell lineage tree by a differentiable vector flow field **v**_*TR*_. The latter was calculated through the temporal averaging of cell displacements by a Gaussian kernel filter (*T* = 10 min) and their spatial regularization. The optimal regularization term resulted in a spatial scale of interpolation of *R* = 20 μm, comprising one or two rows of neighbors depending on the cell size (Supplementary Fig. 2). These operations, in addition to revealing the characteristic length scales of tissue patterning, removed possible artifacts originating from cell tracking errors^22^.

The tensorial analysis was achieved by first calculating the Incremental Deformation Gradient (IDG) tensor field^25–26^ to build an instantaneous time-evolving Eulerian description, i.e. in terms of a spatial reference frame, of the local deformation rate at the level of approximate cell centers. IDG tensor invariants (Supplementary Table 2) defined scalar descriptors of the mesoscopic 3D deformation rate including compression and expansion rates (*P* < 0 and *P* > 0 respectively) and rotation discriminant (*D*). Combining *P* and *D* provided a *τ* index useful to classify the different types of deformations. The distortion rate descriptor (*Q*_*d*_) derived from the deviatoric tensor^3^, quantified the amount of mesoscopic shape changes produced by cell intercalation and/or cell shape changes (Table 1, Supplementary Fig. 3).

**Table 1.** Instantaneous descriptors. Instantaneous deformation measured at one time step is assumed to be infinitesimal. Therefore, linear decomposition of tensors is used. First column: descriptor name. Second column: details of the descriptor values. Third column: interpretation of the descriptor. Scalar descriptors are defined as tensor invariants: *P* for compression and expansion, *D* for rotation discriminant, *τ,* which combines the sign of the previous two descriptors to define topology labels and *Q*_*d*_, which describes the amount of distortion.

|  |  |  |
| --- | --- | --- |
| $P$ | Volume Change Rate<br>$P>0$ Expansion<br>$P<0$ Compression | Expansion and compression of the tissue.<br>Cell nuclei centroids get further away or closer from a centroid taken as reference in consecutive steps |
| $D$ | Rotation Discriminant<br>$>0$ Rotation<br>$<0$ No Rotation | Measurement of rotation compared to linear deformation.<br>The centroids move around a reference centroid significantly more than changing relative distances. |
| $\tau$ | $P$ and $D$<br>1 Exp. Rotation<br>2 Exp. No Rotation<br>3 Comp. Rotation<br>4 Comp. No Rotation | The combination of the signs of $P$ and the $D$ reveals the main topological configurations. |
| $Q_d$ | Distortion rate | Deformation causing tissue rearrangements.<br>The constellation of centroids around a reference centroid changes shape without changing its enclosing volume. |
| $\{e_i\}$ | Strain rate components | Geometrical characterization (eigenvalues and eigenvectors) of the 3D strain rate. |
| $\{d_i\}$ | Distortion rate components | Geometrical characterization (eigenvalues and eigenvectors) of the 3D distortion rate. |

The different descriptors were displayed as spatiotemporal maps with our custom-made visualization interface Mov-IT^8,25^, revealing characteristic patterns of instantaneous mechanical activity in the zebrafish presumptive brain between 6 and 14 hours post fertilization (hpf). The instantaneous descriptor maps provided a new kind of spatiotemporal landmarks characteristic of morphogenetic transitions. At the onset of gastrulation (6 hpf), planar expansion over the yolk cell indicated that tissue shaping in the analyzed spatio-temporal sequences still involved epiboly movements only, with cells moving away from an isotropic source point in the velocity field **v**_*TR*_ (Fig. 1, Supplementary Movie 2–6). As gastrulation proceeded (7-8 hpf), the mechanical activity became dominated by tissue compression at the dorsal side of the embryo. Consequently, a mechanical boundary revealed by the velocity **v**_*TR*_ and the compression and expansion rate *P* emerged between the dorsal and ventral side of the embryo. By mid-gastrulation (8-9 hpf) the distortion rate (*Q*_*d*_) indicated shear along the midline throughout the depth of the tissues, likely to result from the relative movements of hypoblast and epiblast (Fig. 1, Supplementary Fig. 4, Supplementary Movie 7−11). Massive mesoscopic distortion (*Q*_*d*_) and drop of compression observed at the time of tail bud closure (10-11 hpf) was consistent with fast convergence toward the midline and cell intercalation (Supplementary Fig. 4). By 9 to 12 hpf, increasing rotation rates (*D*) on both sides of the midline marked the end of gastrulation and onset of neurulation, with the direction of rotation contributing to tissue convergence toward the midline. By 12 hpf, the disappearance of large scale patterns indicated another regime of tissue dynamics (Fig. 1).

**Figure 1.**
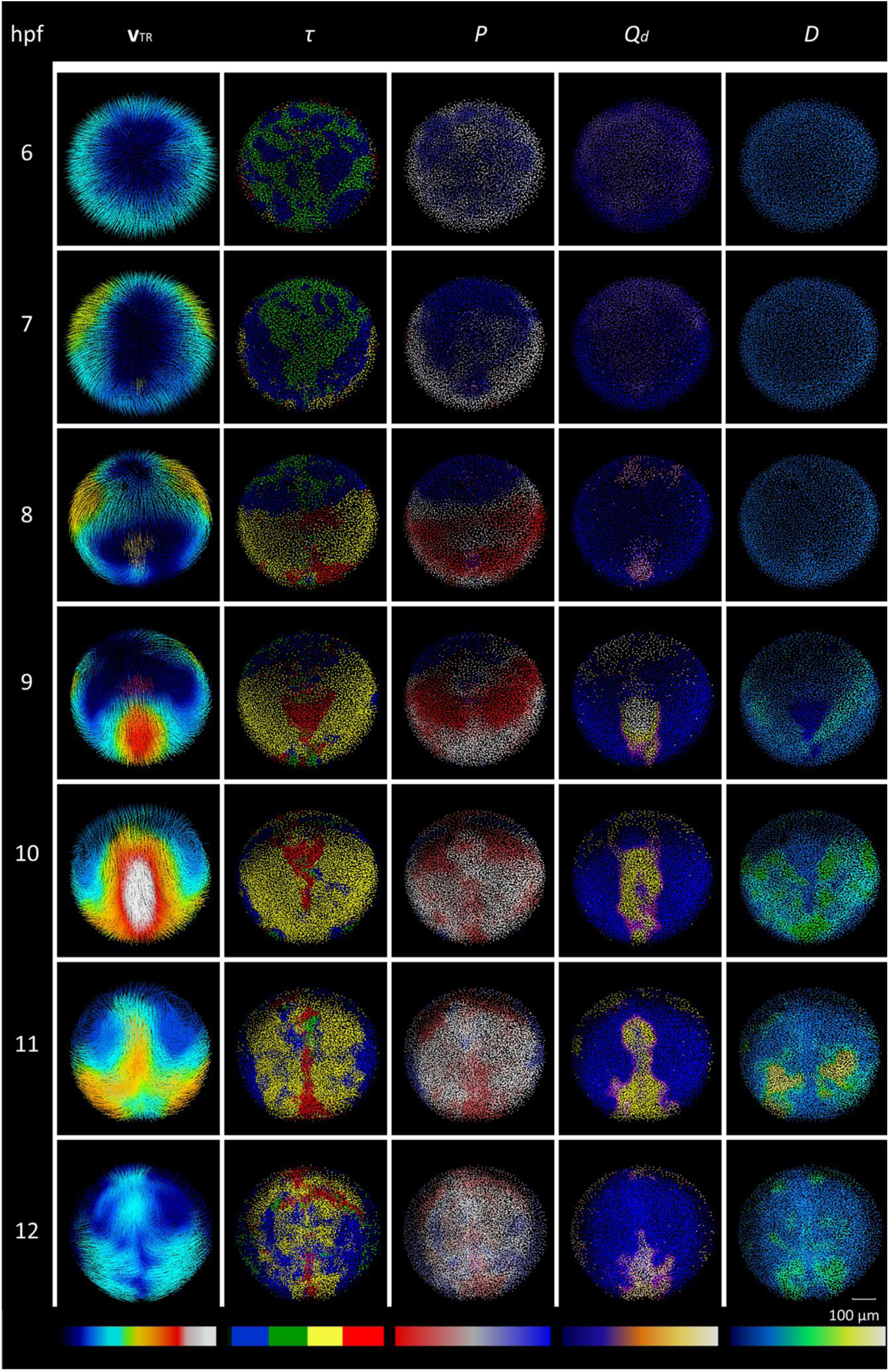
Phenomenology of gastrulation from instantaneous deformation descriptors. Patterns of instantaneous deformation descriptors throughout gastrulation and early neurulation (6-12 hpf) in zebrafish wild-type embryo, animal pole view (z1). Rows correspond to chosen developmental times (hpf). Columns display descriptors (table S2, S3): **v**_*TR*_ (regularized velocity field), *P* (compression/expansion rate), *Q*_*d*_ (distortion rate) and *D* (rotation discriminant) and *τ* (index as the combination of signs of *P* and *D*). Color map at the bottom of each column indicates minimum to maximum values from left to right. Scale bar 100 μm.

The mechanical patterns revealed by the instantaneous descriptors (Fig. 1, Supplementary Movie 2−6) displayed heterogeneities in space and time. While the velocity (**v**_*TR*_) field was at least from 6 to 10 hpf fairly symmetric, the pattern of the topology descriptor (*τ*), combining compression (*P*) and rotation (*D*) measurements and cumulating their fluctuations, displayed bilateral heterogeneity, and its persistence in time was blurred by instantaneous fluctuations. These observations suggested that, while constrained in the tissue flow of gastrulation, neighboring cells experienced, in a 10 min time scale, variable strain levels combining compression, distortion and/or rotation.

### The robustness of biomechanical cues results from the cumulative deformation rates along cell trajectories

We hypothesized that cumulating the mechanical constraints experienced by the cells over finite time intervals could make more robust and persistent patterns unfold^20^ (Fig. 2).Assessing the cumulative deformation rates along cell trajectories relied on a Lagrangian trajectory-based representation of the descriptors defining Lagrangian Biomechanical Profiles (LBPs), which will be further classified and averaged in CLBPs (Supplementary Fig. 5). Computing series of IDG tensors along the trajectories integrated from the vector flow field **v**_*TR*_ (movie S12) provided a sequence of Finite-Time Deformation Gradient (FTDG) tensor fields. Cumulative mechanical cues along the trajectories from an initial temporal reference *t*_ini_ were then expressed by the invariants of the FTDG tensors (Table 2, Supplementary Table 2). The latter included volume changes (*J*), rotation angle (*θ*) and tissue distortion or reshaping (*MIC1* and *MIC2*) for the amount of distortion and its geometrical configuration respectively (Fig. 2, Supplementary Movie 13). The cumulative LBPs led to spatially more homogeneous territories than the Eulerian instantaneous descriptors (Supplementary Movie 13, 14) suggesting, that throughout gastrulation, neighboring cells experienced similar cumulative mechanical deformations (Fig. 2, Supplementary Fig. 6).

**Table 2.**
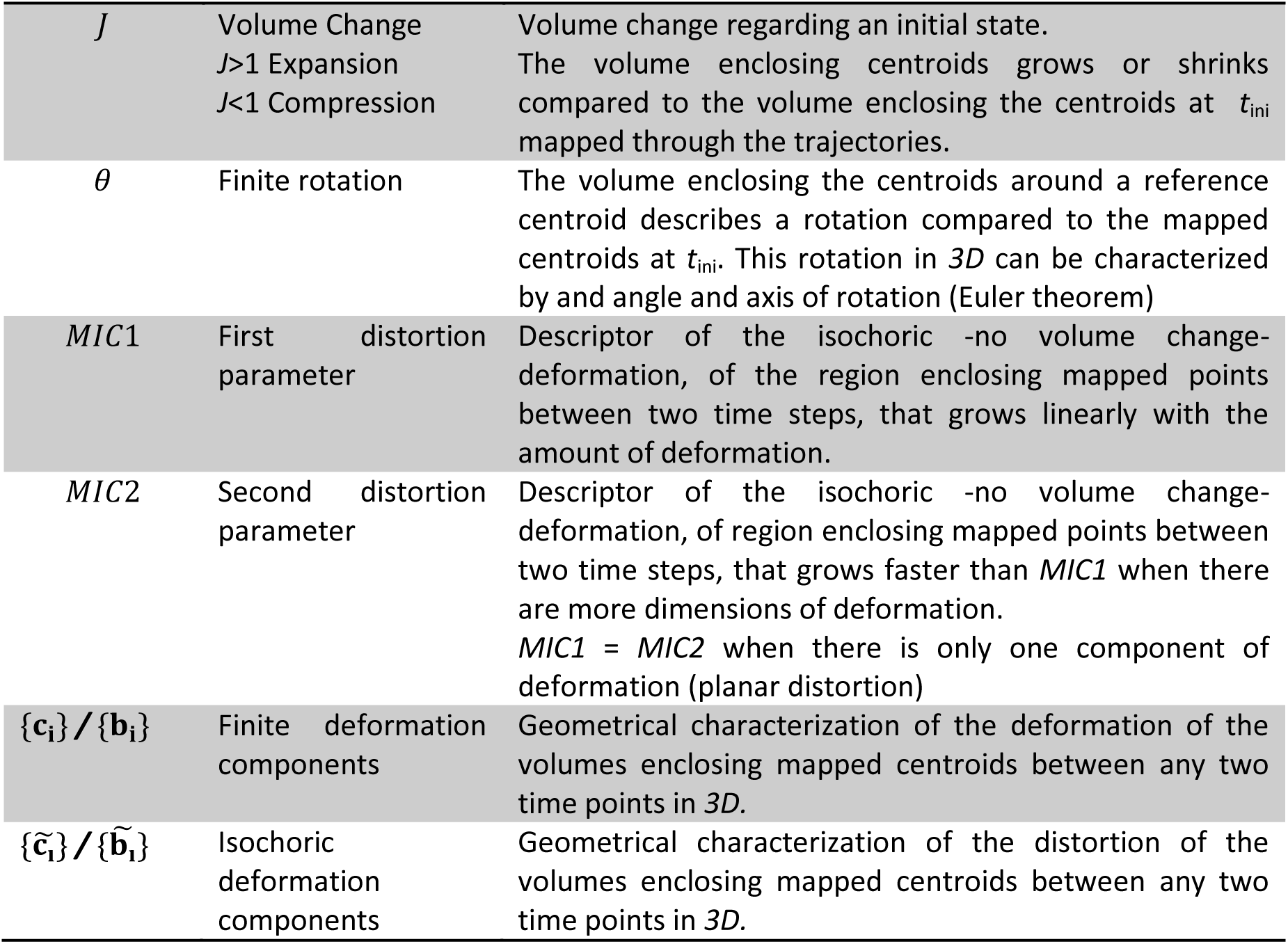
Finite time descriptors. Finite-time deformation descriptors. First column: name of the descriptor. Second column: short name of the descriptor. Third column: interpretation of the descriptor. These descriptors describe deformation through time given an initial reference state: J for compression and expansion, θ for finite rotation around an axis of rotation, MIC1 and MIC2 for distortion of the tissue in 3D and the principal components of strain and isochoric (not producing volume changes) strain.

|  |  |  |
| --- | --- | --- |
| $J$ | Volume Change<br>$J>1$ Expansion<br>$J<1$ Compression | Volume change regarding an initial state.<br>The volume enclosing centroids grows or shrinks compared to the volume enclosing the centroids at $t_{ini}$ mapped through the trajectories. |
| $\theta$ | Finite rotation | The volume enclosing the centroids around a reference centroid describes a rotation compared to the mapped centroids at $t_{ini}$. This rotation in 3D can be characterized by an angle and axis of rotation (Euler theorem) |
| $MIC1$ | First distortion parameter | Descriptor of the isochoric -no volume change- deformation, of the region enclosing mapped points between two time steps, that grows linearly with the amount of deformation. |
| $MIC2$ | Second distortion parameter | Descriptor of the isochoric -no volume change- deformation, of region enclosing mapped points between two time steps, that grows faster than $MIC1$ when there are more dimensions of deformation.<br>$MIC1 = MIC2$ when there is only one component of deformation (planar distortion) |
| $\{\mathbf{c}_i\} / \{\mathbf{b}_i\}$ | Finite deformation components | Geometrical characterization of the deformation of the volumes enclosing mapped centroids between any two time points in 3D. |
| $\{\tilde{\mathbf{c}}_i\} / \{\tilde{\mathbf{b}}_i\}$ | Isochoric deformation components | Geometrical characterization of the distortion of the volumes enclosing mapped centroids between any two time points in 3D. |

**Figure 2.**
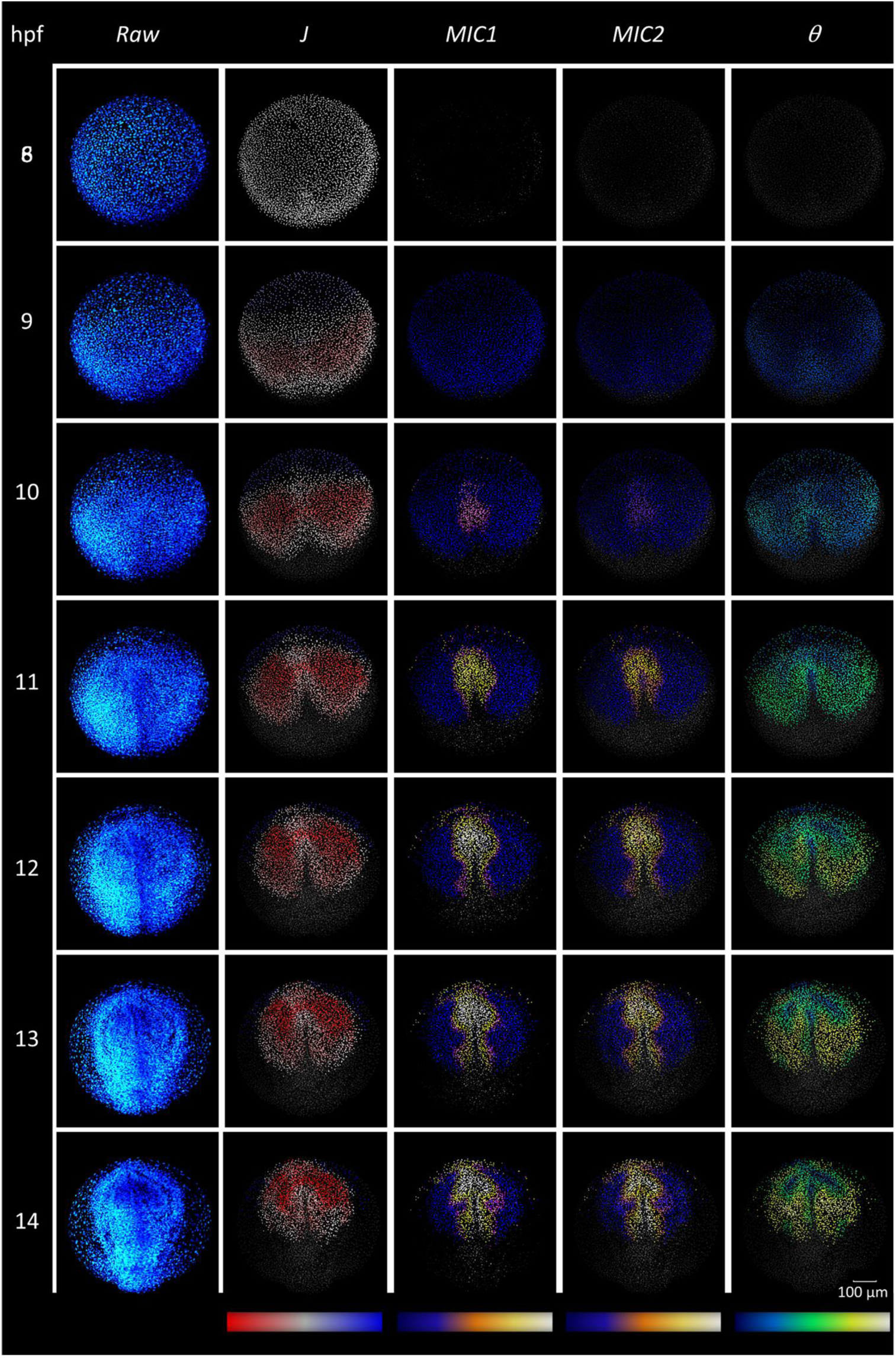
Phenomenology of gastrulation from cumulative deformation descriptors. Same type of display as in Fig. 1 for the embryo z1 with cumulative deformation descriptors from the initial state (*t*_ini_) set at 8 hpf. Rows correspond to chosen developmental times (hpf). Columns display the 3D rendering of the raw data (nuclear staining) and the descriptors (table S2, S4): *J* (compression/expansion), *MIC1* and *MIC2* (distortion), *θ* (finite rotation). Color map at the bottom of each column indicates minimum to maximum values from left to right. Scale bar represents 100 μm.

### Interindividual comparison based on the Lagrangian Biomechanical Profiles of selected cell populations

To further investigate the homogeneity of cumulative versus instantaneous mesoscopic mechanical cues in cell populations and the robustness of the patterns in different zebrafish embryos, we investigated different ways of decreasing the dimensionality of the data. This was achieved by defining cell populations either a priori according to embryological knowledge or without any a priori hypothesis using machine learning methods. The mean and variance of instantaneous LBPs for similar cell domains encompassing part of the dorsal epiblast and hypoblast selected manually in five different embryos^27^, (tailbud selection, Supplementary Fig. 7, Supplementary Movie 15) confirmed temporal markers for the progression of gastrulation in both tissues (Fig. 3A, Supplementary Fig. 4). Interindividual comparison confirmed the robustness of biomechanical features but also indicated timing differences, which we interpreted as a consequence of variability in experimental conditions (e.g. temperature). The onset of epiblast compression was taken as a landmark to temporally align the different datasets at an initial state (*t*_ini_ between 7-8 hpf) and calculate the cumulative profiles (Fig. 3B). As expected from results in Fig. 2, cumulative LBPs produced more homogeneous and robust patterns than the instantaneous ones, reinforcing the hypothesis of spatiotemporal compensation of local fluctuations.

**Figure 3.**
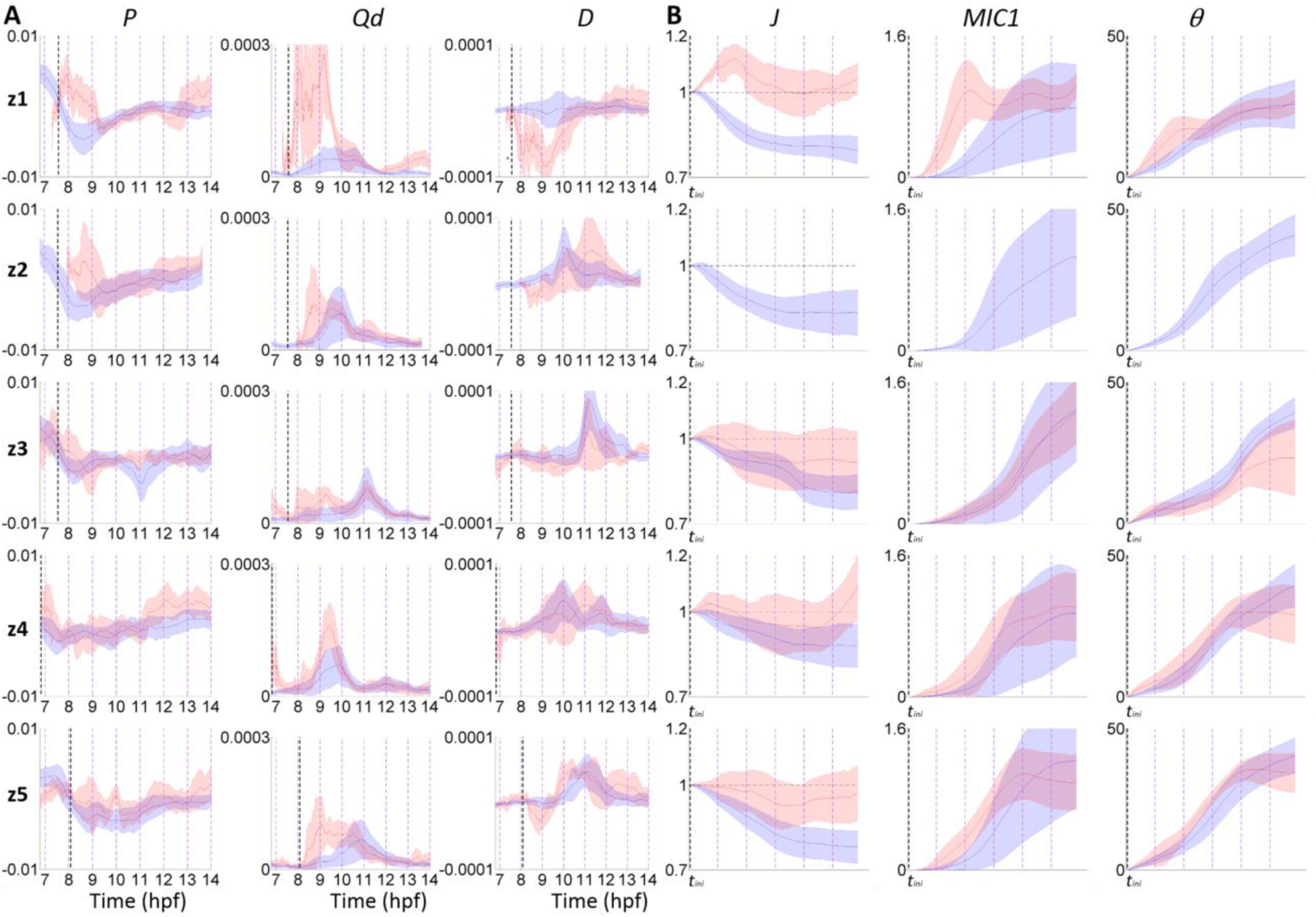
Quantitative comparison of Lagrangian Biomechanical Profiles (LBPs) in a cohort of zebrafish embryos. Comparison of the LBPs’ mean (line) and variance (shaded area), calculated for selected cell populations (tail bud selection, fig. S7, movie S15), hypoblast (red) and epiblast (blue) in embryos z1-z5 (row 1 to 5 respectively). (A) Instantaneous LBPs. Time in hpf. The vertical dashed black line indicates the onset of epiblast compression chosen as reference time (*t*_ini_). (B) LBPs cumulated from (*t*_ini_) for the next 6 hours, mean (line) and variance (shaded area), same cell population (tail bud selection) as in (A). The 5 plots are aligned in time at *t*_ini_. Hypoblast was not analyzed in embryo z2 as it was not present by *t*_ini_ in the cell population selected by the tail bud stage and backtracked (see cell selection fig. S7, movie S15).

### Canonical Lagrangian profiles (CLBPs) to characterize cell population mechanical history

We further investigated the spatiotemporal coherence of the biomechanical patterns through the systematic categorization of the cumulative LBPs for a selection of cells remaining in the field of view from 8 to 14 hpf (shield-stage selection, Fig. 4A, Supplementary Fig. 8). This cell population at the animal pole of the early gastrula was expected to encompass the presumptive forebrain and the underlying prechordal plate^20^. The similarity of the LBPs was estimated based on their dynamic range and temporal evolution. We calculated the LBP distance distribution using a cosine metric and unsupervised classification to find groups minimizing its variance. This classification created categories of LBPs and the corresponding canonical LBPs (CLBPs) for each of the descriptors (Fig. 4A, B). As illustrated in Fig. 4B and Supplementary Fig. 9−S11, three CLBPs were sufficient to characterize the diversity of mesoscopic deformation histories within the selected cell populations. For each of the descriptors, one of the canonical profiles appeared to segregate the hypoblast from the epiblast, indicating its phenotypic homogeneity (Supplementary Fig. 12). Conversely, the epiblast population appeared heterogeneous as two different canonical profiles were necessary to characterize its cells’ mechanical history (Supplementary Fig. 12).

**Figure 4.**
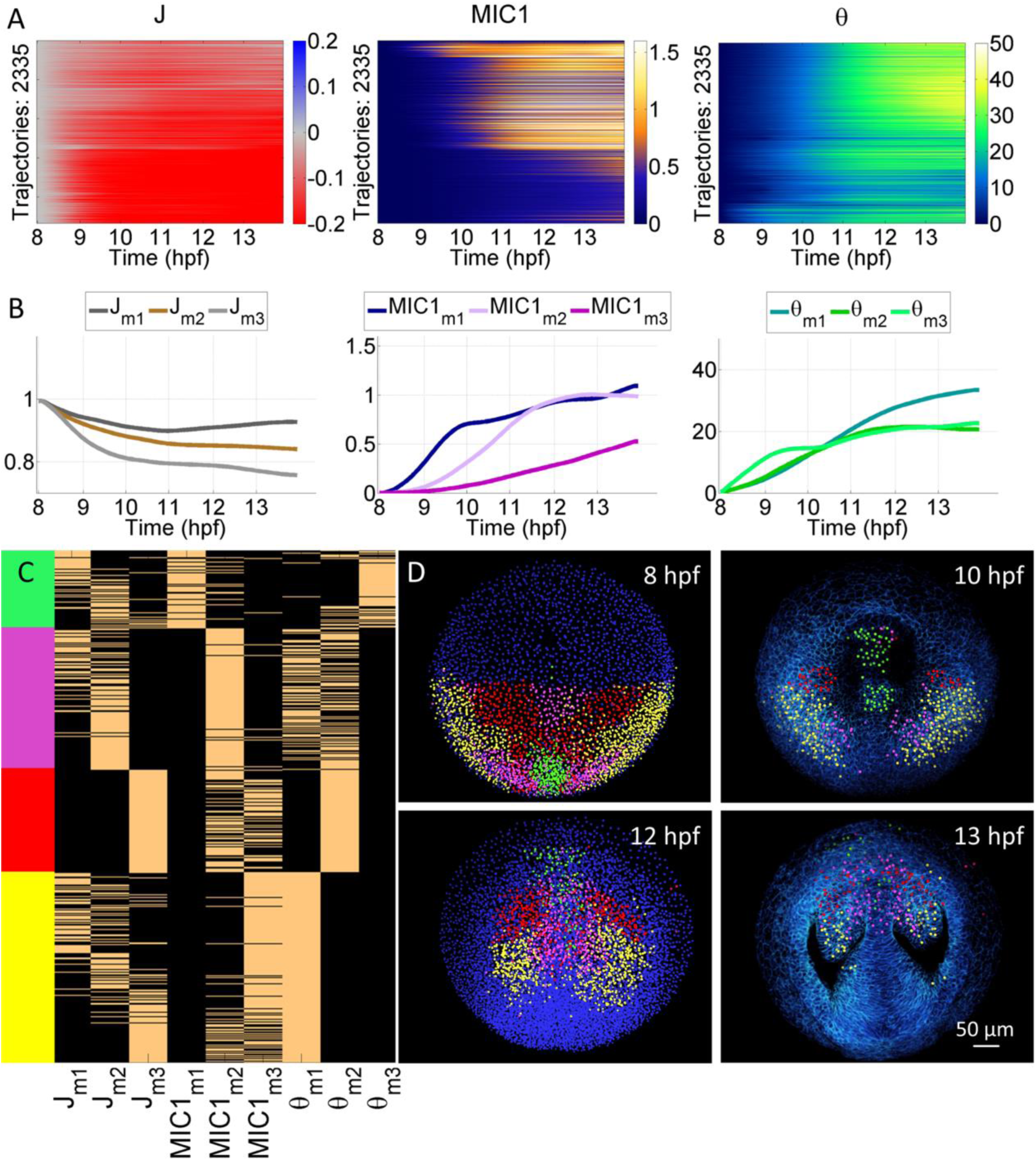
Order and coherence of the Lagrangian biomechanical map. (A) Lagrangian Biomechanical Profiles (LBPs) for selected cells (shield selection, fig. S8). Each panel displays LBPs prior clustering (y-axis) through time (x-axis) for *J*, *MIC1*, *MIC2* and *θ*. (B) The mean (m) for each of the three of LBP clusters (number identified as optimal fig. S9–S12) defined the CLBPs (e.g. *J*_m1_ to *J*_m3_ for *J, etc*). (C) Mechanical signature of each trajectory as a 9-tuple built according to its greatest similarity (1 beige or 0 black) to LBP clusters (fig. S13) (B). Hierarchical clustering of the cells according to their signature led to four clusters (green, red, purple and yellow). (D) The four clusters identified at *t*_ini_= 8 hpf were displayed using Mov-IT and the color code was propagated along the cell lineage (fig. S14, S15, movie S16–S18). Snapshots at 8, 10, 12 and 13 hpf indicated top left. At 8 and 10 hpf, all the detected nuclei are displayed with unlabeled nuclear centers in blue. At 12 and 13 hpf, only the labeled nuclear centers are displayed with 3D rendering of membrane raw data in blue and upper sections removed down to 65 μm below the embryo surface. Scale bar 50 μm.

### Unsupervised classification of cell trajectories according to their CLBPs leads to an ordered and coherent biomechanical map

The temporal and spatial coherence and robustness of the cumulative descriptor patterns suggested that unsupervised machine learning strategies could reveal morphogenetic domains with distinct mechanical histories. The cell trajectories were hierarchically classified according to their signature in terms of CLBPs (Fig. 4C, Supplementary Fig. 13) revealing four main populations spatiotemporally ordered and coherent and forming bilaterally symmetric domains to be compared with the state-of-the-art brain fate map^20,28^ (Fig. 4D, Supplementary Fig. 14, Supplementary Movie 16, 17). We used our interactive visualization tool Mov-IT to compare the CLBP based categories to morphological compartments identified with the cell membrane fluorescent staining. The green domain confined at the posterior midline by 8 hpf, and undergoing an early increase in distortion and rotation followed by a rigid anterior displacement over the yolk, was interpreted as gathering the prechordal plate and the ventral forebrain, i.e. presumptive hypothalamus. The similar mechanical histories of these two fields highlight the role of the prechordal plate in the formation of the hypothalamus and separation of bilateral eyes. The three other populations (yellow, pink and red) matched the eye field and probably the ventral telencephalon. The eye field appeared divided into three domains with different mechanical histories: a most ventral domain (yellow) that underwent by the end of gastrulation the steepest temporal increase in rotation, a ventral medial domain (pink) that underwent a late distortion and low-to-intermediate compression and a ventral lateral (red) subjected during gastrulation and early neurulation to the highest increase in compression (Supplementary Fig. 15, Supplementary Movie 18). Altogether, we propose that the unsupervised classification of canonical LBPs characterizing the cumulative mechanical cues along cell trajectories identifies morphogenetic fields and anticipates their regionalization.

## Discussion

The digital reconstruction of cell lineage trees from 3D+time imaging of developing living systems provides a new kind of data suitable to investigate and model the cell and tissue dynamics underlying morphogenetic processes^29^. The full exploitation of this huge amount of quantitative data requires the development of new methodologies in an interdisciplinary context. State-of-the-art algorithmic workflows designed to process time-lapse imaging data and achieve cell tracking in space and time have limitations and do not produce error-free lineage trees^21,22^, especially in the case of vertebrate organisms with high cell density and thick tissues. A typical 2% error rate of false links per time step requires hand corrections to fully retrieve the cell clonal history. But we demonstrate that this somewhat noisy data is readily amenable to kinematic analysis based on a continuous tissue flow approximation^30^. With their characteristic deformation time scale of 10 min and spatial scale of 20 μm, the zebrafish gastrulating tissues possess a fluidlike behavior, as suggested in other vertebrate species (e.g. chicken^31^). The possibility to work within this paradigm opens interesting theoretical possibilities, although the limits of its application to living cells and tissues have not been fully explored. In particular, a constitutive equation that would accurately capture the spatially and temporally heterogeneous stress/strain relationship within embryonic tissues has yet to be established. To do so would require stress measurements in the whole organism and this kind of experimentation is a major challenge. In this context, our 3D framework for the automated kinematic analysis of tissue deformation provides unique information to unveil the possible role of mechanical cues in morphogenetic processes. Although leaving open the question of forces driving zebrafish gastrulation, our 3D kinematic analysis indicates that in wild-type embryos, the displacement of the hypoblast relative to the epiblast at the dorsal midline generates bilateral jet-vortices that shape the anterior brain including bilateral eyes. The 3D kinematic patterns and the comparison of their spatial and temporal dynamics with the patterns corresponding to different outputs of biological processes such as the fate map, provide a new set of tools to decipher the causalities underlying the formation of morphogenetic fields and presumptive organs. More generally, 3D instantaneous and cumulative tissue deformation patterns, validated here with zebrafish gastrulation, would readily give insights in morphogenetic processes for unknown species. The method is well suited for large-scale collective cell displacements imaged and reconstructed with state-of-the-art strategies.

## Methods

### 3D+time imaging data of a cohort of zebrafish embryos

Wild-type *Danio rerio* (zebrafish) embryos were stained as described by RNA injection at the one-cell stage with 100pg H2B-mCherry and 100pg eGFP-HRAS mRNA (fig. S1) prepared from PCS2+ constructs^32,33^. Embryos raised at 28.5°C for the next 3 hours were dechorionated and mounted in a 3 cm Petri dish filled with embryo medium. To position the embryo, the Petri dish had a glass coverslip bottom, sealing a hole of 0.5mm at the dish center, holding a Teflon tore (ALPHAnov) with a hole of 780 μm. The embryo was maintained and properly oriented by infiltrating around it 0.5% low-melting-point agarose (Sigma) in embryo medium^34^. Temperature in the Petri dish slightly differed for the 5 specimens (z1: about 25°C estimated, z2: 26°C z3: 26°C z4: 28.6°C z5: 24.7°C given by a temperature probe in the Petri dish (OKOLAB). After the imaging procedure, the embryo morphology was checked under the dissecting binocular and the animal was raised for at least 24 hours to assess morphological defects and survival. The different datasets encompassed the same developmental period (4-6 hpf to 14-16 hpf). All the specimens were imaged from the animal pole and the imaged volume encompassed the forebrain with some differences depending on the animal positioning in its mold. Variability in the development speed reflects temperature differences as well as intrinsic variability of embryonic development. Imaging was performed as described^22^ with 2-photon laser scanning^35^ on Leica SP5 upright microscopes and high numerical aperture 20x water dipping lens objectives. Image acquisition parameters are summarized in Supplementary Table 1.

### Digital cell lineages

The 3D+time datasets featured a constant time step *∆t* of approximately 2.5 minutes, defining a discrete time scale for the cohort and a voxel size between 1.2 and 1.4 μm3 (Supplementary Table 1). Digital cell lineages (Supplementary Fig. 1) were obtained through the BioEmergences image processing workflow^22^. The cell lineage data including cell positions at each time step, linkage from one time step to the other and linkage between mother and daughter cells at the time of cell division is presented in a comma-separated values (.csv) table format (cell identifier and position, mother identifier called as such whether the cell divides or not). The BioEmergences tracking method provided an error rate of approximately 2% representing the percentage of false or missing links between two consecutive time steps (*t* to *t+∆t*). Cell positions were given by the approximate nucleus centers.

### Flow field approximation of cell lineage

We considered the cell lineage to provide complete structured spatiotemporal information about cell trajectories, including cell divisions. Specifically, we proposed a generalized data record for each cell nucleus within the lineage as follows:

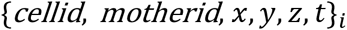

where *i* indexes the detected cells, *cellid is* their corresponding unique identifier at the (*x,y,z,t*) spatio-temporal nucleus position and *motherid* is the identifier of the linked nucleus position at the previous time step (mother cell). The set of detected nuclei forms a spatiotemporal discrete map {**x**(*x*, *y*, *z*, *t*)}_*i*_ of the imaged embryo. Given each detected nucleus at position **x**_*i*_ and time step *t*, the displacement field **v**(**x**_*i*_) was calculated by searching the corresponding *cellid* positions at *t+∆t*. In case of a dividing cell, two matches were found, and the one minimizing the Euclidean distance of the displacement was chosen. The vector field was then generated iteratively from the displacements between matched nuclear positions.

Given a temporal resolution of the time-lapse data of *∆t* ~ 2.5 minutes, singular cell displacements produced by divisions or tracking errors occurred at a frequency close to the sampling’s Nyquist frequency and were thus assumed to generate high-frequency noise. In order to filter out this noise, we performed a temporal smoothing of the displacements along the cell lineage with a Gaussian kernel *N*(0,*T*), where *T* is a scale in the order of minutes. By testing several parameters, we set *T =* 10 min to generate the smoothed displacement field **v**_*T*_(**x**_*i*_). In addition, displacements over the threshold *MaxMov =* 9 μm/∆*t* were removed as outliers.

We performed a regularization of the cell displacement field to obtain a continuous vector flow field representing the best differentiable approximation of the cell displacements. The vector field **v**_*T*_ was interpolated around the position of each detected nucleus **x**_*i*_ in order to be locally differentiable using a Gaussian kernel *N*(0, *R*):

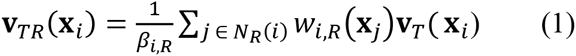

where 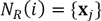 is the set of neighbors interpolated, 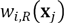 is the weight of each neighbor according to the *N*(0, *R*) distribution and *β*_*i,R*_ the sum of all the weights. In order to preserve boundaries within the displacement field, a binary function 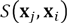 was used to discard outlier displacements:

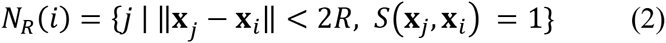

where 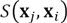 comprises three thresholds based on data observation: the maximum angle of deviation against the reference (*π*/2), a minimum speed (0.2 μm/min) and a maximum ratio of speed against the reference 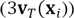. We determined the lengthscale of the regularization filter by balancing the spatial resolution and differentiability of the resulting vector flow field **v**_*TR*_, as described below.

The vector field 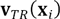 represents a discrete inhomogeneously sampled field. To quantify the differentiability of this field, we used a local second-order structure function in terms of the average velocity differences within *n* concentric rings around each **x**_*i*_. Local differentiability is observed when the structure function follows a power law with exponent *γ ≥* 2. The structure function is defined as:

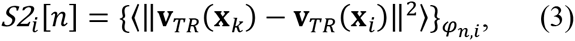

where *n* ∈ [1,10] and *φ*_*n,i*_ denotes that we calculated the discretized function *S2*_*i*_[*n*] around each position **x**_*i*_ in concentric rings with the same radius *dl*, i.e.

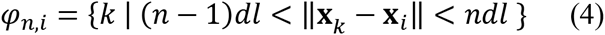

We determined the regularized vector field **v**_*TR*_ and the kernel width *R* considering the local differentiability of all field samples. To this end, we computed a time-dependent ensemble average of the *S2*_*i*_[*n*] function for each ring 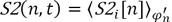, where 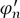 is the subset of all the rings derived from *φ*_*n,i*_ after removal of outlier rings. We labeled a ring as an outlier when the function 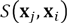 described above for the cells within the ring was negative for the majority of them, indicating largely divergent information in the ring that, if included in the regularization term, could over smooth the field globally. Testing several parameters, an optimal kernel width was found, *R =* 20 μm, that ensured differentiability while minimally worsening the spatial resolution of the displacement field (Supplementary Fig. 2).

### Instantaneous deformation descriptors

The differentiability of the vector flow field **v**_*TR*_ allowed us to apply principles of continuum mechanics to quantify cell motion and tissue deformation. We calculated the Incremental Deformation Gradient -IDG- tensor field **f**(**x**_*i*_) by a numerical method (least squares error minimization) considering the displacements of the neighboring cells within the volume defined by *2R.* The IDG tensor for the cell *i* at position 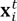 and time *t* defines a mapping from the material vector 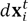 onto the vector 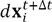:

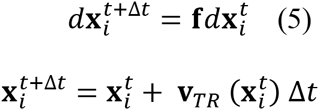

We obtained flow topology descriptors as Galilean invariants derived from the tensor Incremental Gradient of Displacements **h** = **f** – **I** (Table 1, Fig. 1). These invariants are tensor metrics that are independent of the reference frame orientation and velocity. They are therefore suitable for visualizing and comparing complex 3D flows. The first, second and third principal invariants in *3D* (referred here as invariants) are interpreted as follows. The first invariant of **h** (*P* as described in 26) quantifies the compression/expansion rate at the mesoscopic scale, corresponding to an overall decrease/increase of cell size change in the vicinity of each nucleus. The second invariant (*Q* as described in 26) gives information about deformation, not producing volume changes at the mesoscopic scale, associated with both irrotational and vortical motions. We identified the deformation produced by rotation the discriminant of the deformation tensor (*D*), which is positive in regions of mesoscopic rotation. Finally, we designed a topology index descriptor *τ* that takes four different values representing the combinations of the signs of the descriptors *P* and *D* (Fig. 1): expansion-rotation (green label), expansion-no rotation (blue label), compression-rotation (yellow label) and compression-no rotation (red label).

To further characterize the strain rates, we calculated the symmetric part of the tensor **h** that stands for the irrotational, incremental strain tensor **ε** and its principal components. This symmetric tensor generally provides information about shears and changes in volume through its second invariant *Q*_*s*_. For tissues that may change volume and in order to distinguish between reconfigurations of the tissues (cell intercalation and cell shape changes) and volume changes (cell size), we calculated the deviatoric tensor **d** that subtracts volume changes from the strain rate. The eigenvectors {**d**_1_, **d**_2_, **d**_3_} and second invariant *Q*_*d*_ of this tensor provided information on the tissue distortion associated to collective cell intercalation and cell shape changes (Table 2, Fig. 1, Supplementary Fig. 3, 4).

### Building Lagrangian Biomechanical Profiles

We defined a Lagrangian representation of the flow field by approximating the reconstructed cell trajectories by the flow path lines of the regularized vector flow field. Lagrangian analysis of non-compressible (*P*=0) two-dimensional flows have been successfully applied to discover Lagrangian Coherent Structures in fluid transport^36–38^. Here, we proposed Lagrangian metrics based on the computation of finite-time deformation tensors to unfold the biomechanical history along the lineage. Because of cell divisions and incomplete cell trajectories, the reconstructed cell lineage had to be regularized to build a continuous flow description. We interpolated the cell trajectories with the information of the flow field displacements to generate complete trajectories given an interval of time [*t*_*n*_, *t*_*m*_] (Supplementary Movie 12). Thus, we generated a bijective spatio-temporal map 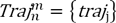 suitable to express the dynamics in terms of trajectories (Lagrangian) 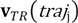 instead of spatial points (Eulerian).

We built Lagrangian Biomechanical Profiles (LBPs) independent of spatial coordinates using the trajectory flow field. LBPs expressed the instantaneous and cumulative biomechanical activity along each trajectory (Supplementary Fig. 5). The cumulative activity was computed for each trajectory by setting a temporal reference *t*_ini_ and by incrementally enlarging the interval of analysis to generate a sequence of FTDG tensors 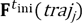 (varying along the time interval of the trajectory) and their corresponding descriptor fields. The computation of the FTDG tensor and its invariants (Table 2) for each interval [*t*_ini,_ *t*] t is described below.

### Computation of FTDG tensors and descriptors

The IDG tensor field was expressed in Lagrangian terms using the trajectory field **f**(*traj*_*j*_) or 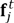. We composed the tensors along each trajectory and the corresponding time interval with the chain rule (forward-projection matrix operation) to generate a Finite Time Deformation Gradient (FTDG):

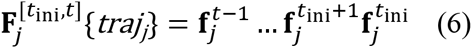

The third invariant of the tensor (*J* as described in 25) characterized the volume change during the time interval. The finite rotation (tensor **R**) was segregated from elongation (tensor **U**) through a polar decomposition **F** = **RU**^39^. The rotation was described with the angle of rotation ***θ*** and the axis of rotation (Euler’s theorem as described in 25):

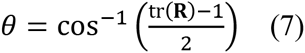

The strains were obtained with the right Cauchy-Green tensor **C** = **F^T^F** and the left Cauchy-Green tensor **B** = **FF^T^** and their principal components. The invariants of these tensors integrated volume changes and shear strains. Therefore, we calculated the isochoric deformation tensor and the corresponding isochoric Cauchy-Green tensors (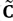 and 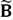) to identify the distortion along the trajectories from the volumetric changes:

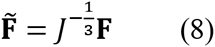

The first and second 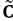 and 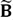 (*MIC1* and *MIC2* as described in 25) represented the tissue shape changes along the time intervals. The descriptor *MIC1* evolved linearly with the amount of distortion whereas the descriptor *MIC2* added second-order terms when the distortion had more than one dimension. Thus together, the *MIC1* and *MIC2* descriptors characterized the amount and geometry of tissue shape changes (Table 2, Fig. 2, Supplementary Fig. 6).

### Visualization of descriptor maps and manual selection of cell domains

The BioEmergences custom visualization tool Mov-IT^22^ was used to explore the 3D+time descriptor maps. The maps for the Eulerian descriptors were computed by generating a color map for the IDG tensor values at each nuclear center **x**_*i*_. For the cumulative LBPs, color maps were built and visualized with the Mov-IT software by assigning values to the closest nuclear center **x**_*i*_.

Mov-IT was also used to manually select cell domains and propagate the selections along the cell tracking, in order to perform a statistical analysis of the corresponding LBPs. Two different types of cell populations were selected. Expert embryologists selected cell populations at 10-11 hpf within the hypoblast and epiblast layers that were approximately similar in position and cell number between the five specimens of the cohort. These populations were backtracked to identify the corresponding progenitors at the onset of gastrulation (tailbud selection Fig. 3, Supplementary Fig. 7, Supplementary Movie 15). We also selected in embryo z1 by the onset of gastrulation the largest possible population of cells kept into the imaged volume throughout the whole imaging sequence (shield selection, Fig. 4, Supplementary Fig. 8, Supplementary Movie 16, 17). This selection was used to categorize the different types of profiles with unsupervised classification.

### Categorization of Lagrangian Biomechanical Profiles

A trajectory field defined from the shield cell selection (Supplementary Fig. 8) was further characterized by identifying subdomains with similar LBPs 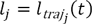 (descriptor along the trajectories). The subdomains were identified by generating a distance *d*_LBP_ distribution between the LBPs (*l*_*j*_, *l*_*k*_) of pairs of cells for each descriptor with a cosine metric, selected because it properly weighted both the magnitude of the descriptor and its deviations along time:

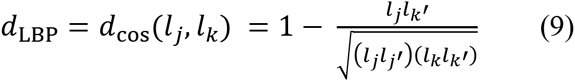

We applied unsupervised *k*-means clustering to classify the trajectories minimizing the variance of the distance distribution, so that trajectories with similar profiles according to the metric were classified together. The behavior of each cluster was defined using the mean of the trajectory profiles along time, which is considered suitable because the variance was minimized. Several values of the number of clusters *k* were tested (Supplementary Fig. 9, 10), finding that 3 clusters provided a suitable representation of Canonical Lagrangian Biomechanical Profiles (CLBPs) (Fig. 4, Supplementary Fig. 11, 12).

### Identifying morphogenetic domains from their mechanical signature

We generated a mechanical signature *ζ*_*j*_({CLBP}) for each trajectory as a binary feature vector based on the corresponding set of CLBPs (Fig. 4, Supplementary Fig. 13). Trajectories were then compared using the Euclidean distance between their mechanical signatures and classified into four representative domains using hierarchical clustering.

The obtained classification was used to label the nuclei at the onset of gastrulation (*t*_ini_), generating Lagrangian Biomechanical Maps (Fig. 4). The spatio-temporal evolution of the mechanical domains was visualized with Mov-IT by propagating the corresponding labels along the cell trajectories (Fig. 4, Supplementary Fig. 14, 15, Supplementary Movie16−18).

### Code availability

Raw image and cell lineage data are available on the BioEmergences website: http://bioemergences.eu/kinematics/ login: kinematics password: AGBYeL4y All code is available upon request or already available at the BioEmergeces workflow^22^.

## Acknowledgments

We thank René Doursat for critical reading of the manuscript and the BioEmergences laboratory, especially Louise Duloquin and Dimitri Fabrèges for their collaboration and support. This work was supported in part by TEC2013-48251-C2-2-R and IPT-2012-0401-300000 for A.S. and M.L.C., by grants NSF CBET – 1055697 and NIH R01 GM084227 for J-C.A. and by grants EC NEST “Measuring the Impossible” 28892, ZF-Health EC project HEALTH-F4-2010-242048, ANR BioSys Morphoscale, France BioImaging infrastructure ANR-10-INBS-04 and InterDIM 2011 Région Paris Ile-de-France for N.P. and P.B.

## Author contributions

DP developed the framework, designed visualization and analysis, integrated analysis and biological relevance and co-wrote the manuscript. BL designed and developed the framework foundation. TS developed the visualization and co-wrote the manuscript. AB carried out biological experimentation and imaging. JG supervised the mechanics framework. AS co-funded and co-coordinated the project. PB co-funded and co-coordinated the project and designed the framework foundation. JCA co-funded and co-coordinated the project, supervised the mechanics framework and co-wrote the manuscript. NP co-funded and co-coordinated the project, supervised data acquisition and analysis, led the biological interpretation, co-wrote the manuscript. MJL co-funded and co-coordinated the project, led the design of the analysis framework, co-wrote the manuscript.

---

**Supplementary Fig. 1.**
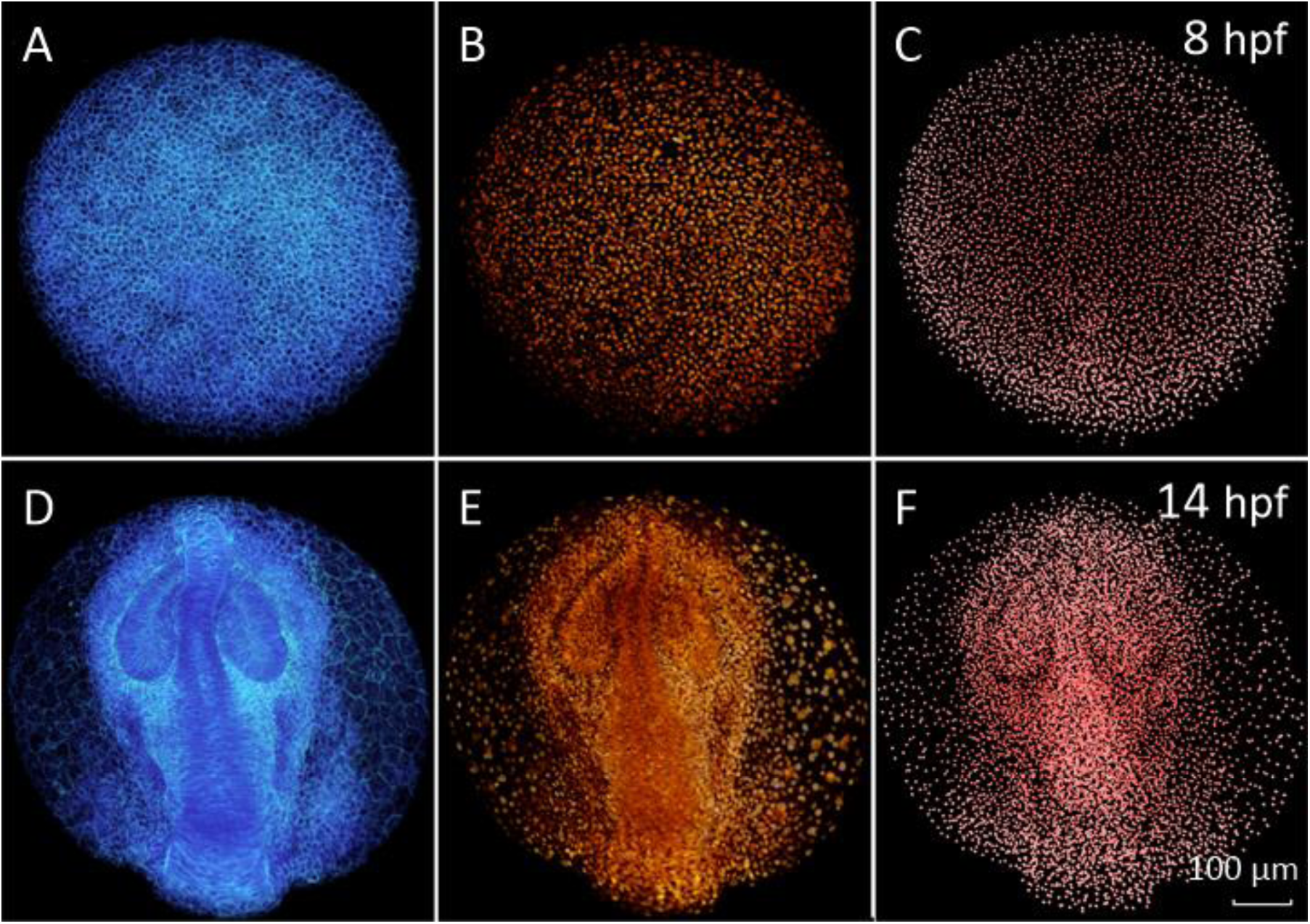
From 3D+time imaging data to digital zebrafish embryo. Wild-type zebrafish embryo, animal pole view, anterior (ventral) to the top, dataset z1 (snapshots taken with Mov-IT). Data acquisition parameters for all datasets z1-z5 are summarized in Supplementary Table 1. (A-C) 8 hpf, (D-F) 14 hpf. (A, D) raw data, membrane staining 3D rendering. (B, E) raw data, nuclear staining, 3D rendering. (C, F) detected nuclei **x**_*i*_.

**Supplementary Fig. 2.**
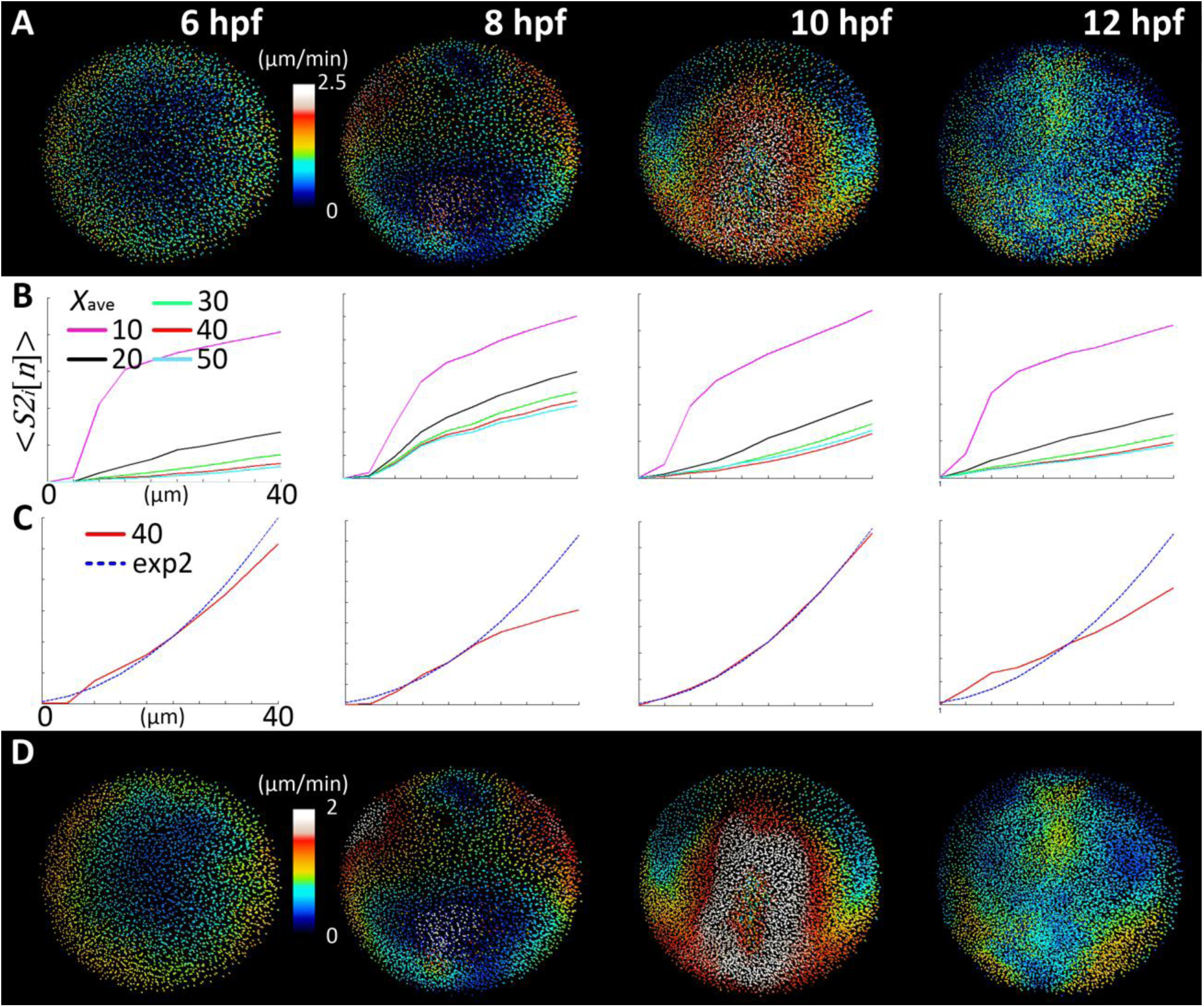
Cell displacement regularization. (A) Displacement field **v**_*T*_(**x**_*i*_) implicit in the cell lineage after applying a temporal smoothing of the displacements along the lineage branches by the Gaussian kernel *N*(0, *T*), (*T* = 10 *mins*), displayed here at different time steps of the development {6,8,10,12} hpf. Colormap goes from zero speed (dark blue) to maximum speed (white). (B) Ensemble-average of the second-order structure function < *S2*_*i*_[*n*] > for all **x**_*i*_ in each time step, plotted as a function of the distance to **x**_*i*_ discretized as *n* concentric spatial rings (*dl* = 4 μm, *n* ∈ [1,10]). Same time steps as in (A). Each curve represents the statistical differentiability given by the ensemble-average at each time step from the regularization of the vector field interpolated with a Gaussian kernel *N*(0, *R*) for *X*_ave_ *=* 2*R =* {10,20,30,40,50} μ*m*. (C) Fitting of the ensemble-average of < *S2*_*i*_[*n*] > function for *2R* = 40 μm to the power law curve of exponent 2 (exp2) that determines the limit of differentiability of the field. Best fit is obtained for the time step 10 hpf. For 12 hpf, the differentiable vector field had a worse fit with the differentiability limit curve. (D) Vector flow field **v**_*TR*_(**x**_*i*_) interpolating displacements using a Gaussian Kernel *N*(0,20). This parameter value preserves the topology of the original vector field **v**_*T*_(**x**_*i*_), but with smoothened and clearer speed information. We interpret that this corresponds to the characteristic lengthscale of the tissue dynamics.

**Supplementary Fig. 3.**
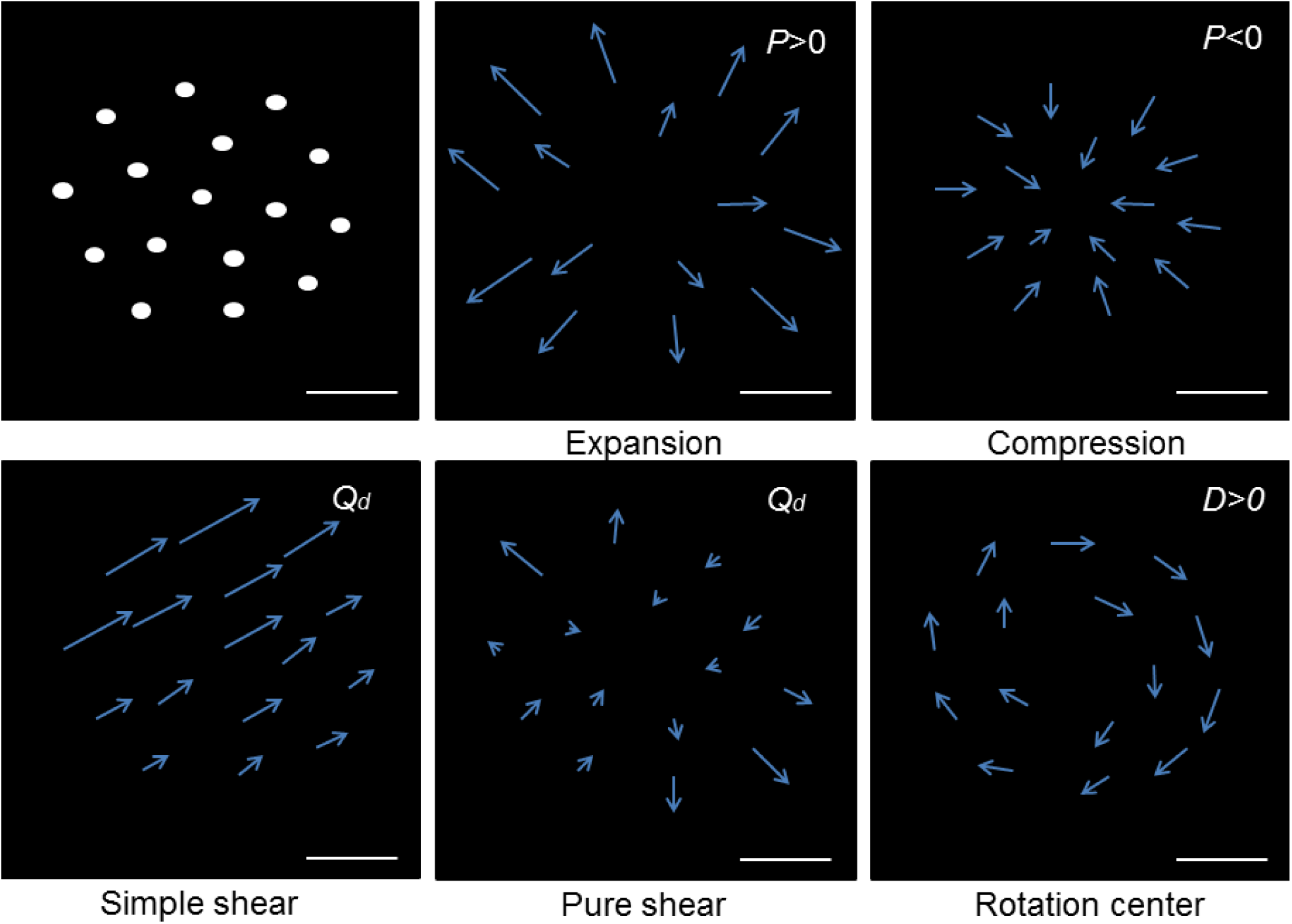
Schemetic representation of instantaneous descriptors. Synthetic examples of instantaneous deformation patterns quantified with the *IDG* tensor and the instantaneous descriptors (Supplementary Table 3). Top-left shows an initial distribution of nucleus centers. The approximate scale bar represents the mesoscopic spatial scale *R=*20 μm (between 1 and 2 neighbors), calculated as explained in Supplementary Fig. 2. The different panels show topologies of the displacements fields characterized by the *IDG* tensor and the derived invariants. Expansion corresponds to growing divergent displacements from a given spatial reference (*P*>0). Compression is characterized by convergent displacements (*P*<0). At high *Q*_*d*_ values, anisotropy of displacement magnitude along a direction generates simple shear and anisotropy of the displacements from a given reference point generates pure shear. Displacements spinning around a reference generate a rotation center and high *D* positive values.

**Supplementary Fig. 4.**
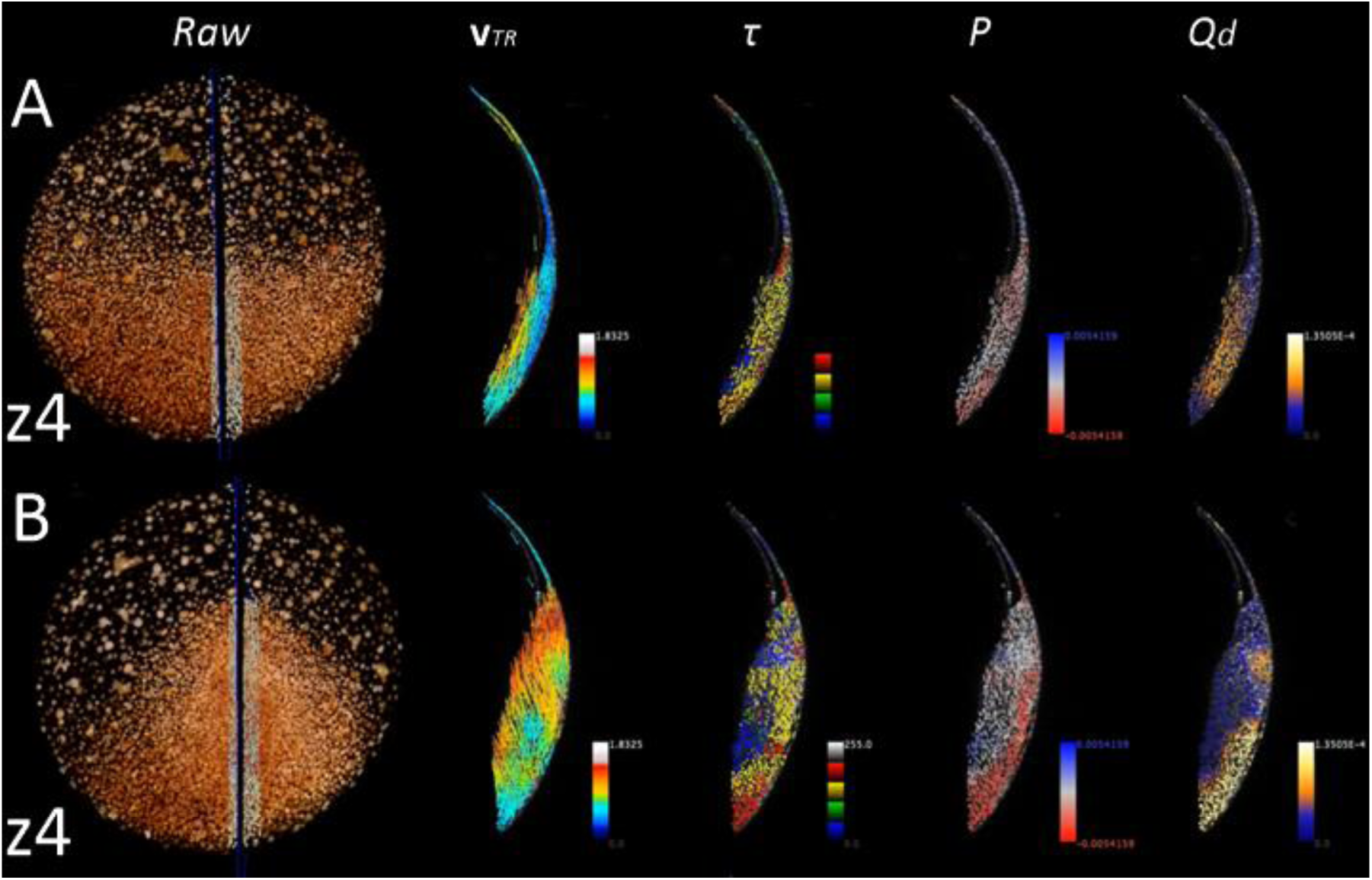
Distortion of cell layers at the midline during zebrafish gastrulation. Instantaneous descriptors 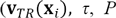 and *Q*_*d*_) along the midline. (A,B) Left panel: 3D rendering of the raw data (raw), animal pole view with a black line at the level of the midline. Selection of cells on both sides of the midline (in white) observed in lateral view in the four other panels. Each descriptor (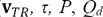 indicated top of each column) is displayed with its color code (bottom right of each panel). (A) 8:45 hpf (embryo z4). We interpret that the descriptors distinguish the hypoblast from the epiblast as the hypoblast has a higher **v**_*TR*_. The relative movements of the hypoblast and epiblast produce simple shear across the depth of the tissue as shown by *Q*_*d*_. The hypoblast compresses less than the epiblast as shown by *P* and *τ*. (B) as in (A) by 10:30 hpf. The epiblast undergoes compression *(P)* along the midline as well as high distortion (*Q*_*d*_) indicating tissue rearrangement, likely to be a consequence of cell intercalation.

**Supplementary Fig. 5.**
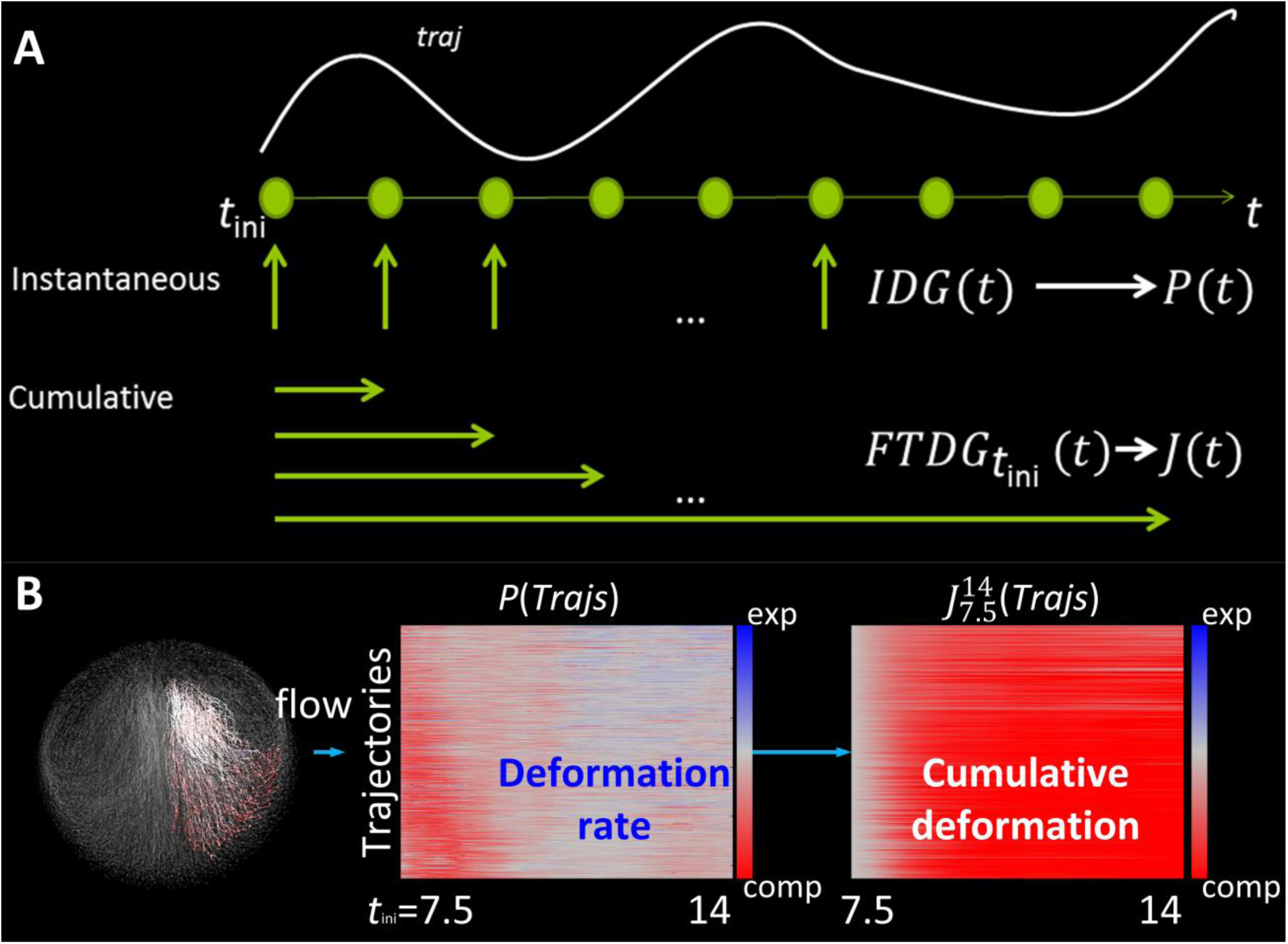
Lagrangian Biomechanical Profiles. (A) Schematic representation of the computation of instantaneous and cumulative descriptors from the *IDG* tensor field expressed in terms of the trajectories (e.g. Lagrangian representation of the *IDG* tensor field). (B) Schematic representation of instantaneous (middle panel) and cumulative (right panel) Lagrangian Biomechanical Profiles (LPBs) for a selected cell population (dataset z1) from 7.5 to 14 hpf. The cell lineage for that selection is colored with the descriptor P and displayed with Mov-IT (left panel). The *P*-LBP (middle panel) shows the compression (red) - expansion (blue) rate history through time (7.5-14 hpf). The *J*-LBP (right panel) shows the cumulative compression (red) - expansion (blue) history from the initial state *t*_ini_ = 7.5 hpf, calculated by applying the scheme described in (A) to each trajectory.

**Supplementary Fig. 6.**
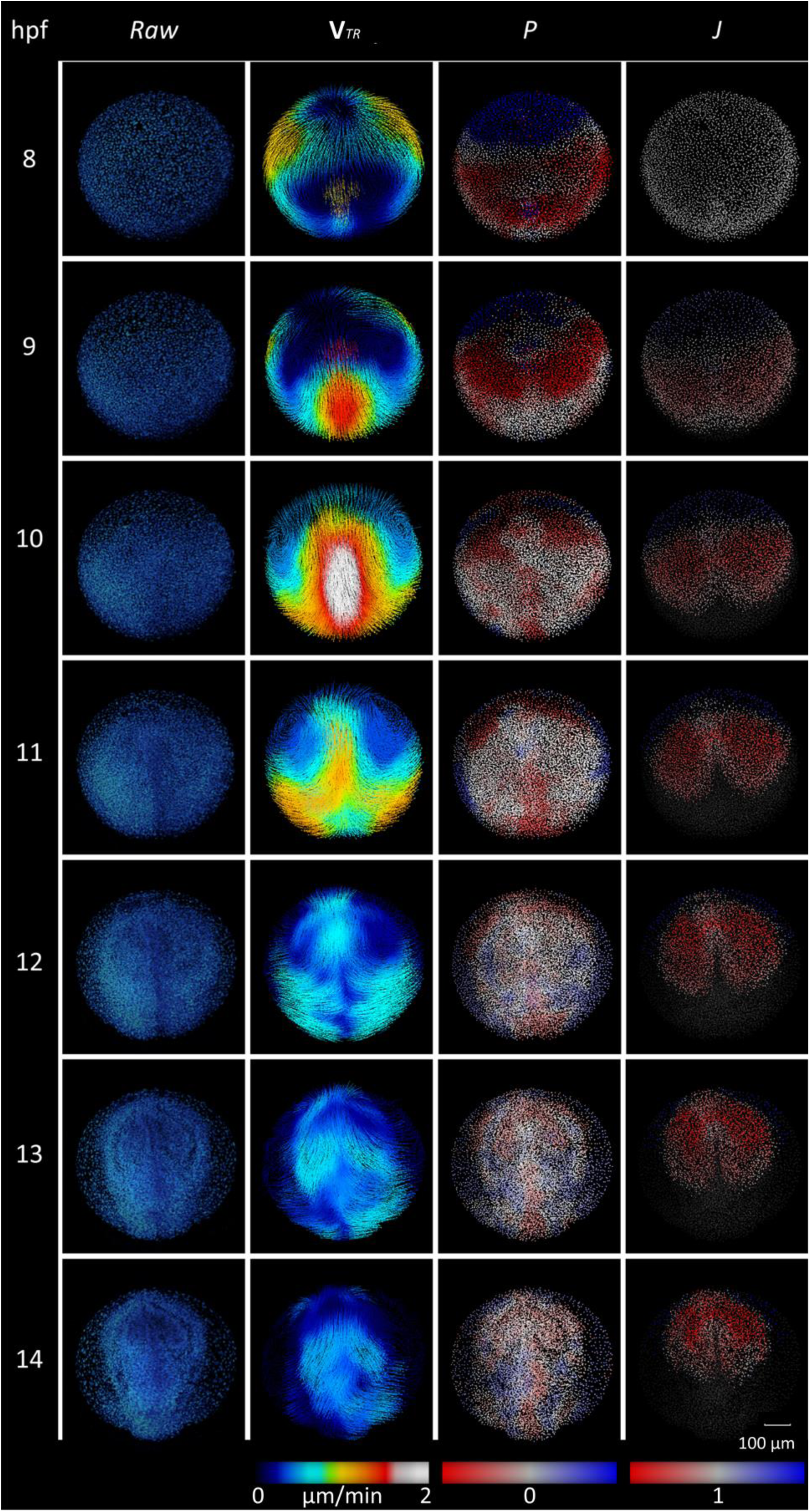
Instantaneous *vs* cumulative compression and expansion descriptors. Same type of display as in Fig. 1, 2. First column: raw data, 3D rendering of nuclear staining. Second column: **v**_*TR*_(**x**_*i*_) as in Fig. 1. The color map from zero speed (dark blue) to the maximum of speed (white) as in Fig. 1 & Supplementary Fig. 4. Third Column: expansion activity denoted by *P* (Eulerian descriptor). The color map for *P* goes from neutral (white=0) to compression (red) and expansion (blue) as in Fig. 1, S4. Fourth column: cumulative compression denoted by *J* (LBP calculated using *t*_*ini*_ = 8 hpf). Same color map as for *P*, no volume change has value 1 as in Fig. 2. By 12 hpf *P* shows mainly neutral values while maximum compression has been reached for *J*. This observation is consistent with the decrease in **v**_*TR*_ (2^nd^ column), observed by the end of gastrulation (11 hpf), meaning that if tissue deformation happens at this stage, its scale is smaller than the mesoscopic scale defined by our smoothing step (see above).

**Supplementary Fig. 7.**
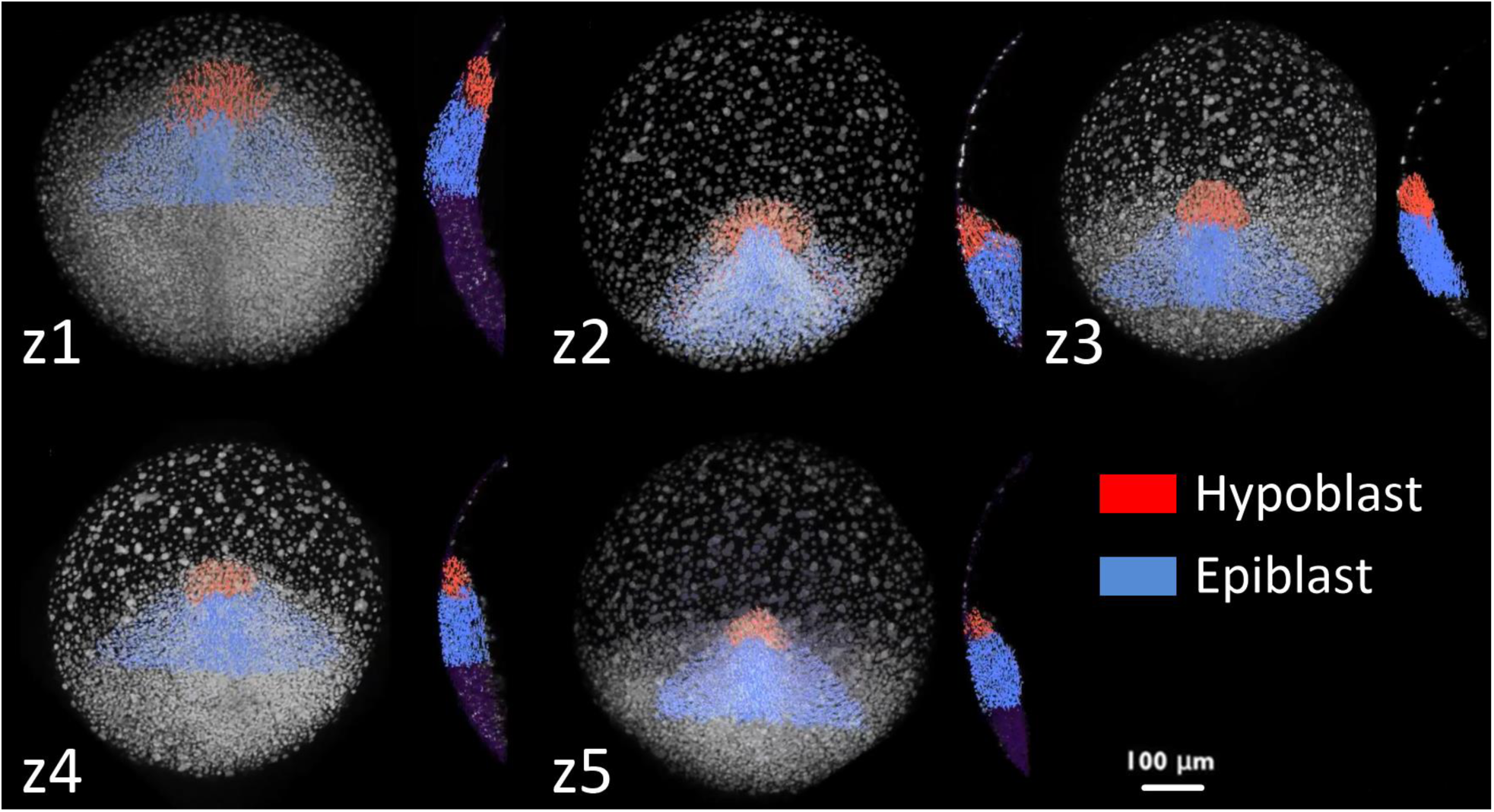
Selection at tailbud of cell populations within the hypoblast and epiblast layers. Embryos z1 to z5 indicated top left. For each embryo animal pole view and lateral view of a sagittal section are displayed, anterior to the top. Snapshots with Mov-IT software display 3D rendering of nuclear raw data and selected nuclear centers. Scale bar 100 μm. Cell populations of similar shape and size were selected in the different embryos by the tailbud stage (10-11hpf). We called this selection “tailbud selection”. Based on the identification of the morphological border separating hypoblast (red) and hypoblast (blue), the two populations were distinguished. These selections are backtracked to define spatio-temporal domains throughout gastrulation (Supplementary Movie 15).

**Supplementary Fig. 8.**
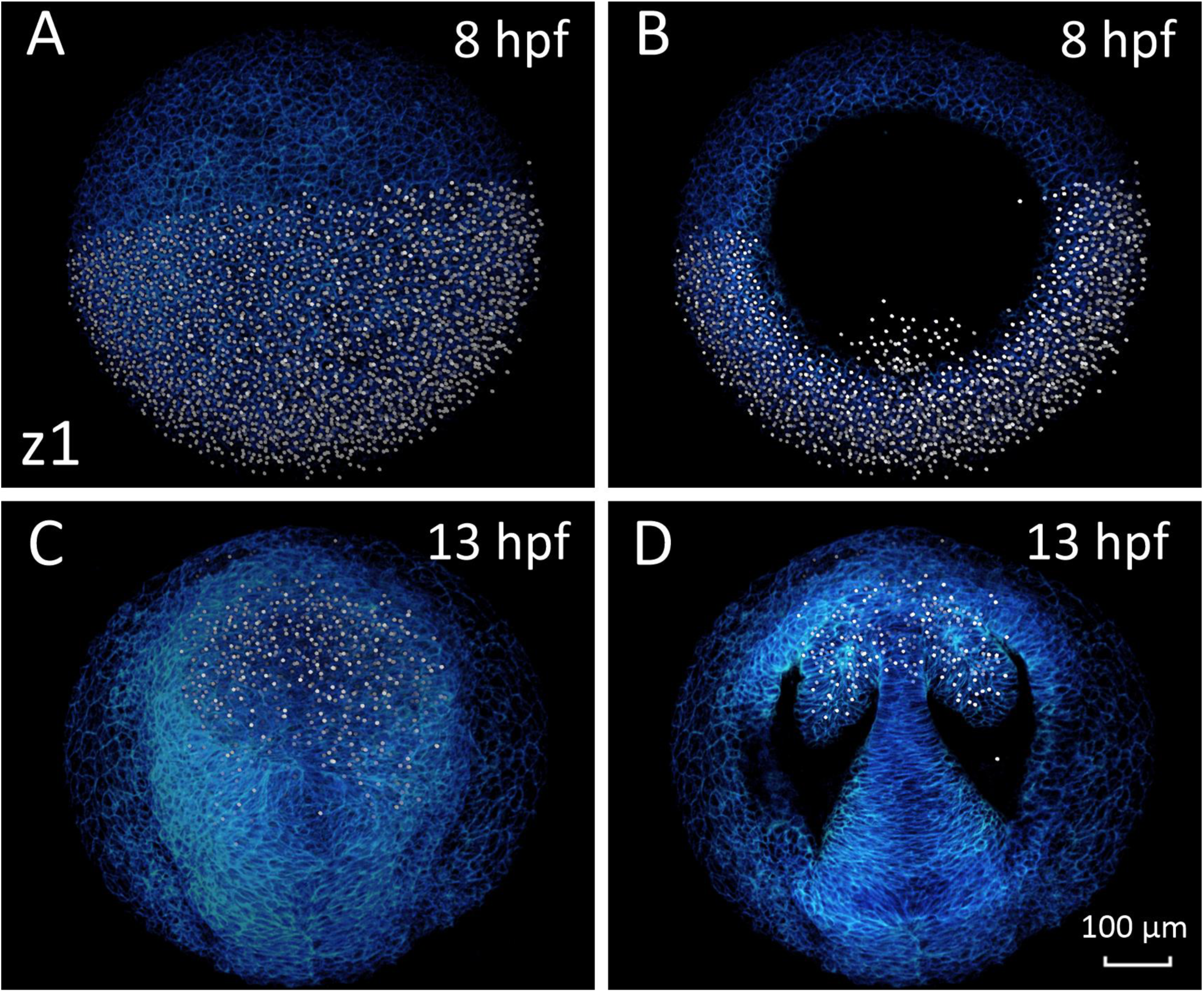
Selection of presumptive forebrain at shield for the unsupervised classification of LBPs. Embyo z1 animal pole view anterior to the top. 8 or 13 hpf indicated top right. Scale bar 100μm. (A) 3D rendering of the whole stack of images at 8 hpf. The domain manually selected with Mov-IT corresponds to the posterior half of the imaged volume through the whole depth of the blastoderm comprising hypoblast cells present the imaged volume (right panel 8 hpf). (B) Same selection but upper embryo sections removed up to 74 μm of depth. This selection was called “shield selection”. (C) 3D rendering of the whole stack at 13 hpf. The selection made by 8 hpf has been propagated through the cell lineage until 13 hpf showing that selected cells are fated to the forebrain including eye vesicles and that the selected cells are kept in the imaged volume. (D) Selection at 13 hpf and upper layers removed at the same depth than B showing a cross-section of the eye vesicles. This selection contains the germ layers selection of z1 shown in Supplementary Fig. 7.

**Supplementary Fig. 9.**
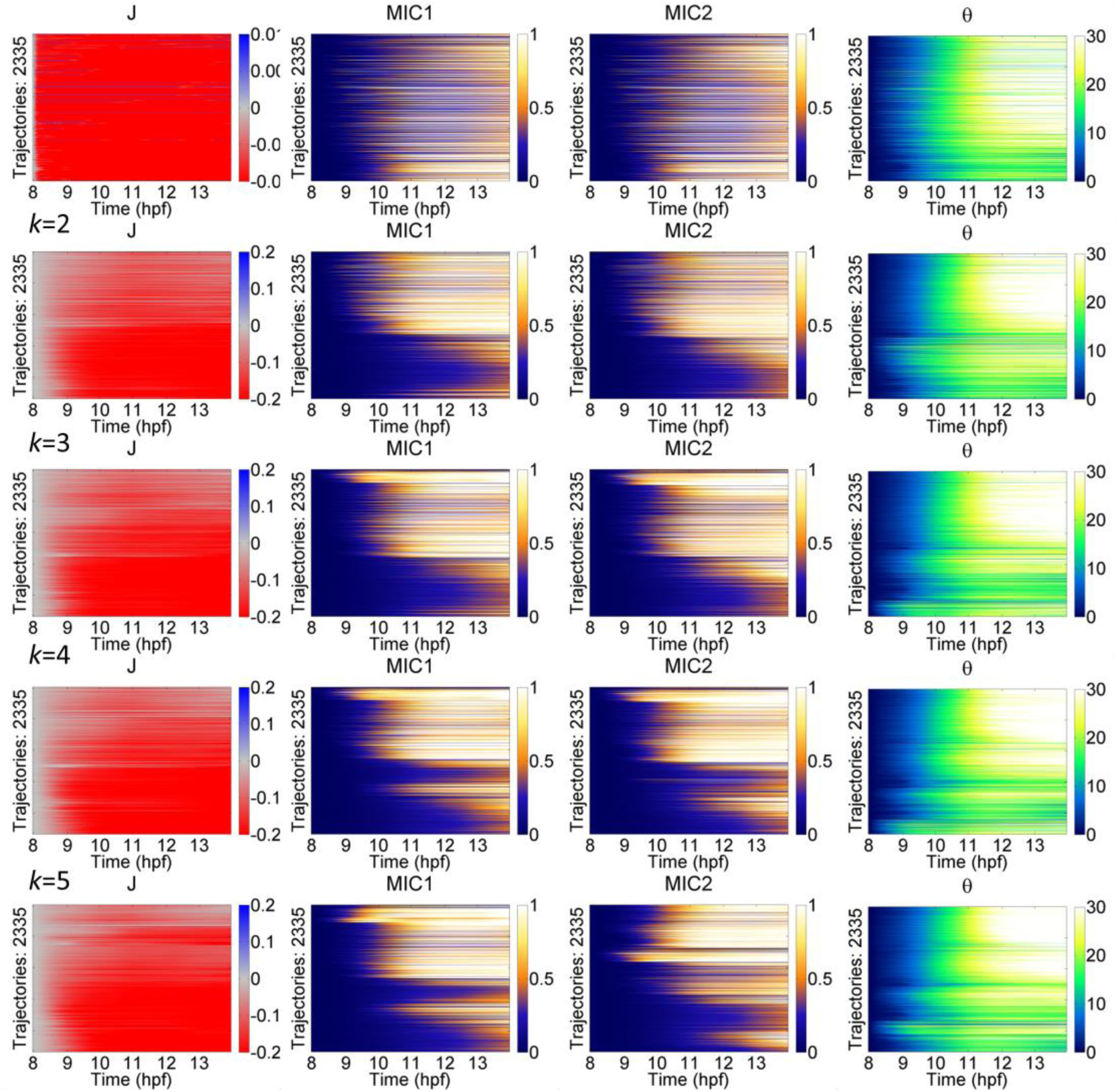
Categorization of Lagrangian Biomechanical Profiles. Row 1: Lagrangian Biomechanical Profiles (LBPs) along the trajectories (ordinates) as a function of time (abscissa in hpf) for cumulative deformation descriptors (columns: *J*, *MIC1*, *MIC2*, *θ*) for the shield selection in embryo z1 (described in Supplementary Fig. 8). Row 2 to 5: *k*-means unsupervised clustering with increasing number of clusters *k* (from 2 to 5). Clustering minimizes the variance of the cosine distance distribution within groups of trajectories. In other words, LBPs are classified and ordered according to their similarity. Colormaps: *J* varies from neutral (white) to compression (red) or expansion (blue). *MIC1*, *MIC2*, *θ* vary from null (dark) to maximum values (white).

**Supplementary Fig. 10.**
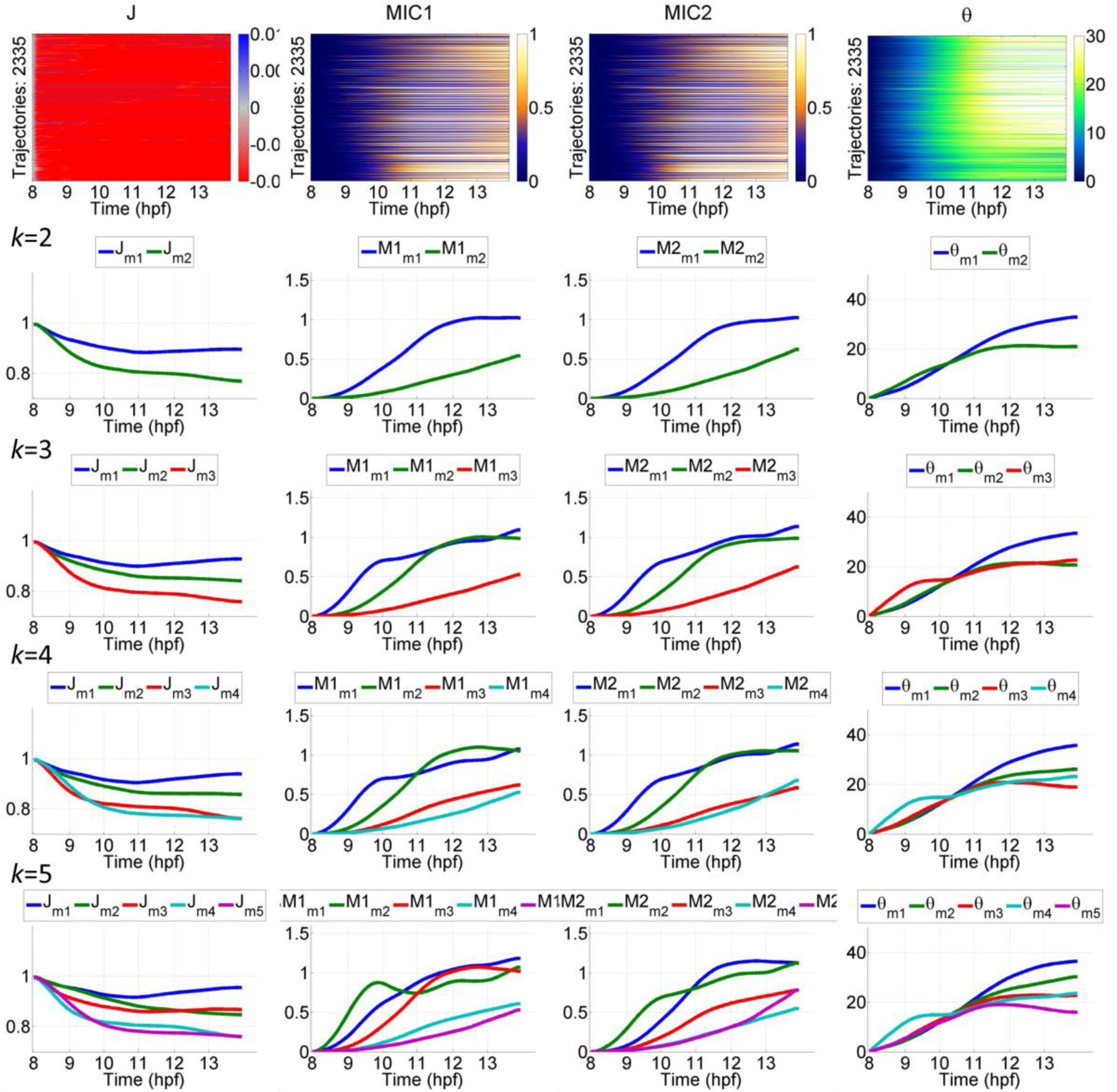
Selecting the relevant number of Canonical Lagrangian Biomechanical Profiles. Row 1: LBPs for each cumulative deformation descriptor (*J*, *MIC1*, *MIC2*, *θ*) for the shield selection (Supplementary Fig. 8) and including epiblast and hypoblast germ layers in the tailbud selection (Supplementary Fig. 7). Row 2 to 5: *k*-means clustering with different *k* parameters (as in Supplementary Fig. 8). The clustering minimizes the variance of the cosine distance distribution of the trajectories. The Canonical LBP is then computed as the mean of the trajectories for each LBP cluster. We observed and validated statistically that increasing *k*>3 did not add significant information to the set of CLBPs.

**Supplementary Fig. 11.**
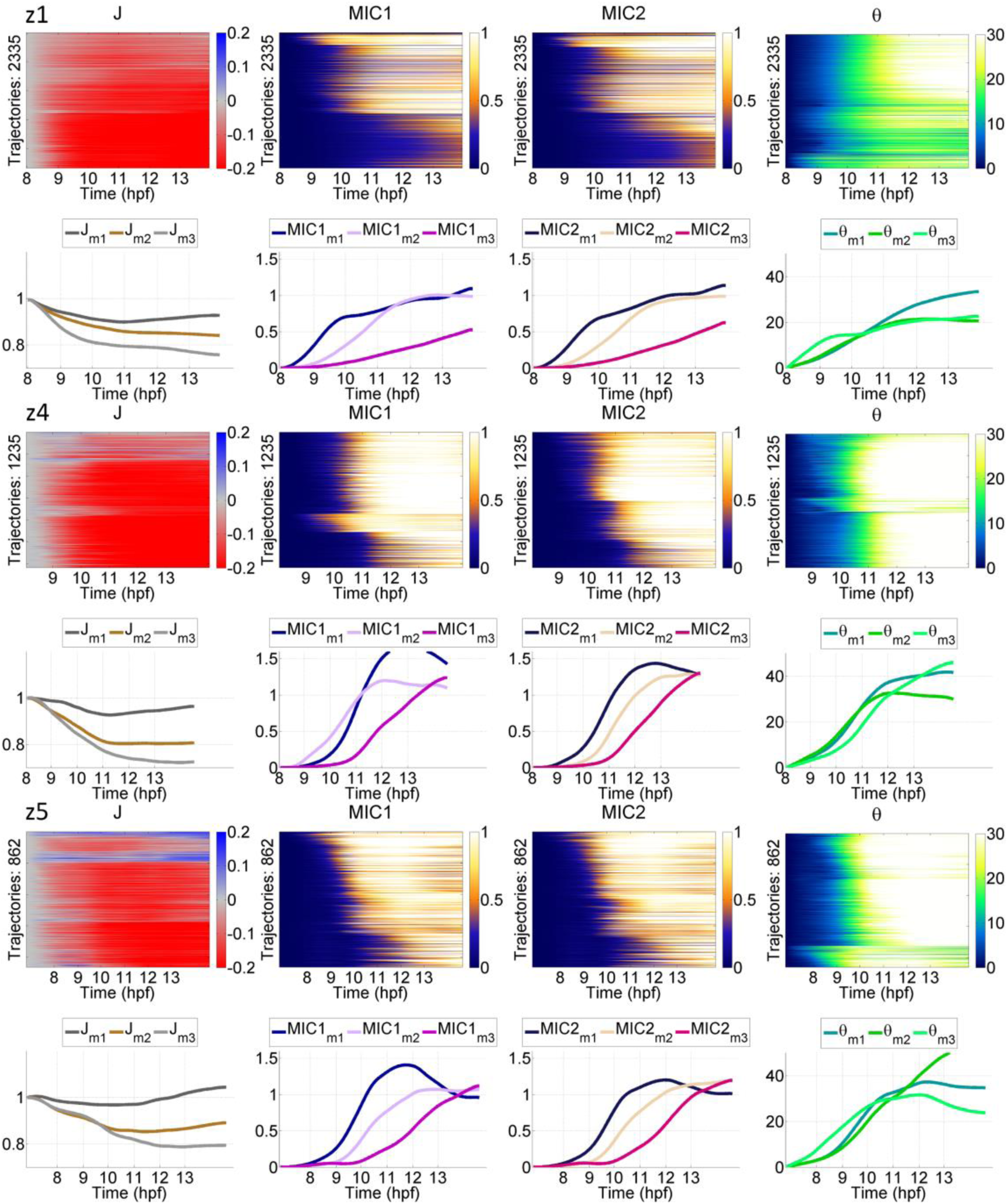
Variability of Canonical Lagrangian Biomechanical Profiles in the Cohort. LBPs for the cumulative deformation descriptors (*J*, *MIC1*, *MIC2*, *θ*) for the shield selection (as in Supplementary Fig. 8) in embryos z1, z4 and z5 (indicated top left of odd rows) corresponding to similar domains comprising epiblast and hypoblast layers. Even rows: plot as a function of time of the corresponding canonical LBPs obtained by *k*-means clustering with *k*=3.

**Supplementary Fig. 12.**
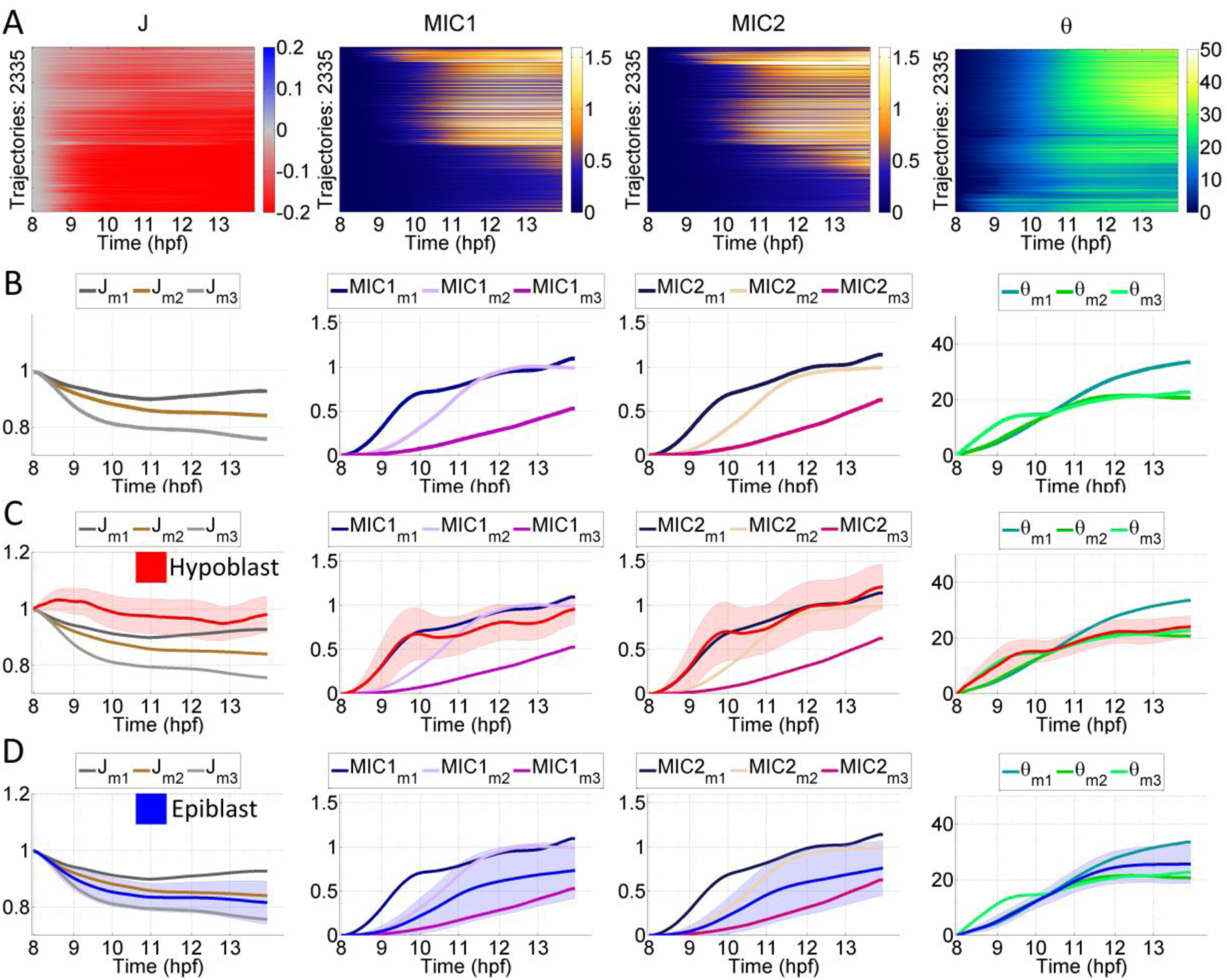
Germ layers (hypoblast and epiblast) and cumulative CLBPs. (A) Lagrangian Biomechanical Profiles (LBPs) of selected cells in embryo z1 (shield selection described in Supplementary Fig. 8). Each image comprises all the LBPs by trajectory (y-axis) along time (x-axis) for *J*, *MIC1*, *MIC2* and *θ*. Trajectories are ordered by comparing their profiles with a cosine metric and using k-means clustering to group trajectories minimizing the variance of the metric. (B) Canonical LBPs for each descriptor as the mean profile of the trajectories within each cluster in (i.e. *J*_*m1*_ for the first CLBP of *J*). Categorization was estimated as optimal with three CLBPs (Supplementary Fig. 9–S11). (C) Average LBP of the hypoblast (red as shown in Supplementary Fig. 7 and Supplementary Movie 15) overlaid to the CLBPs. (D) Average LBP epiblast LBP (blue as shown in Supplementary Fig. 7 and Supplementary Movie 15) overlaid to CLBPs. Two CLBPs were necessary to describe the profile of the epiblast layer.

**Supplementary Fig. 13.**
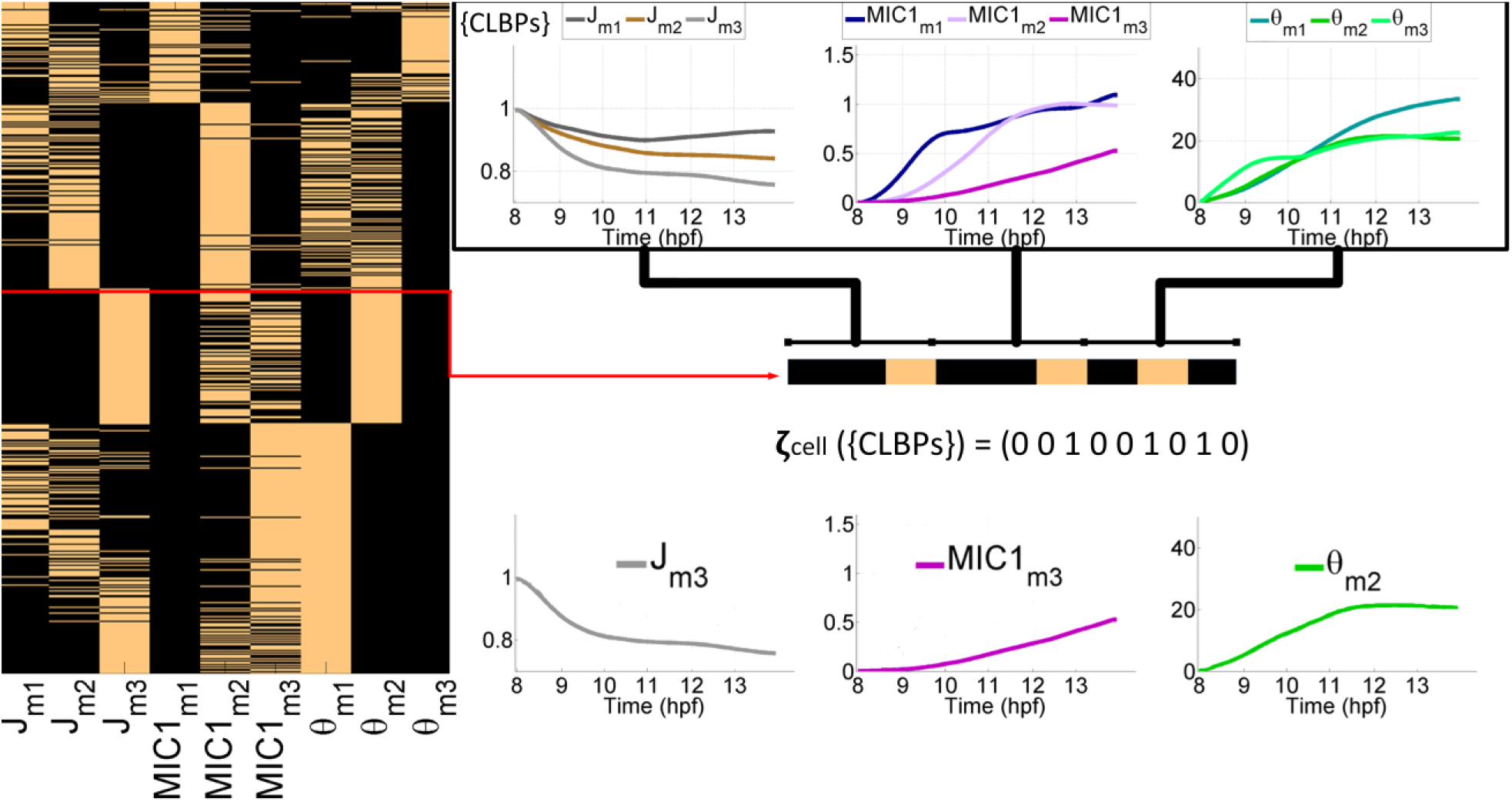
The mechanical signature of cell trajectories. Left panel: Mechanical signatures of trajectories *ζ*_*j*_({CLBP}) for the cells in the shield selection as shown in Supplementary Fig. 8. The mechanical signature of each trajectory (ordinates) is defined by their specific set of CLBPs (abscissa). Right panel: construction of the mechanical signature of a cell trajectory. The signature is a 9-tuple composed of values 0 (black) or 1 (beige) depending on trajectory LBP greatest similarity (1 to the most similar and 0 to the others) to the canonical LBPs of the descriptors (*J, MIC1, θ* shown in the top row). Example of a trajectory with signature (001001010, middle row) resulting in a signature characterized by CLBPs *J*_*m3*_, *MIC1*_*m3*_ and *θ*_*m2*_ (displayed right panel, bottom row).

**Supplementary Fig. 14.**
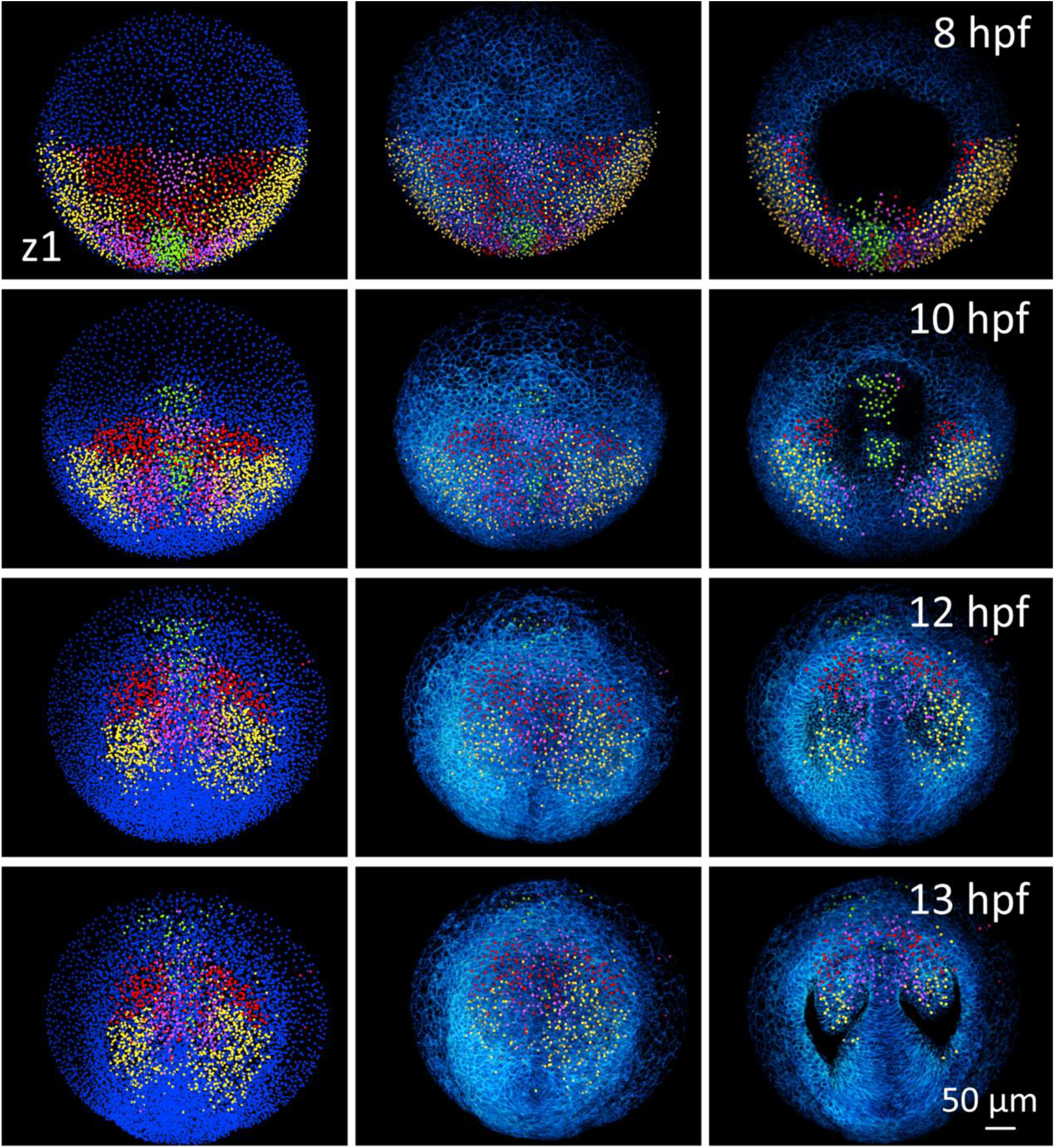
Lagrangian Biomechanical Map and morphological compartments. Lagrangian Biomechanical Maps derived from the categorization of cells in 4 domains by 8 hpf (=*t*_ini_) according to their mechanical signatures. Snapshots by 8, 10, 12 and 13 hpf (top left of each row). Animal pole view. First column: detected nuclear centers with staining of the four domains (green, red, yellow and pink). Second column: nuclear centers in the four domains with their respective colors and 3D rendering of the raw data membrane channel. Third column: same as second column but with removal of upper sections (down to 64 μm below the embryo surface) to reveal morphological landmarks for the hypoblast (8 and 10 hpf) and the regionalization of the eye vesicles (12 and 13 hpf).

**Supplementary Fig. 15.**
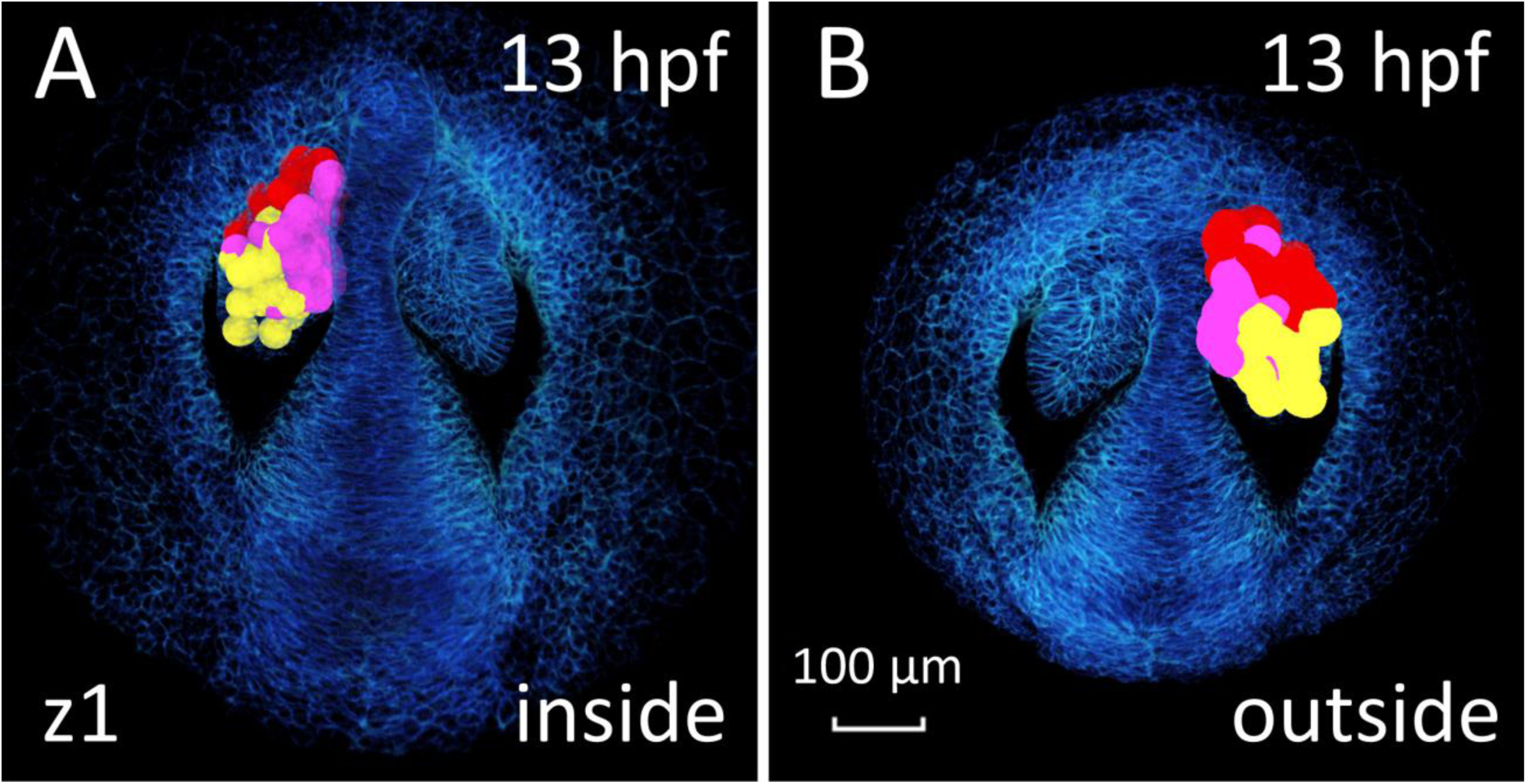
Lagrangian Biomechanical Map of the eye vesicle. Highlights the biomechanical domains in the eye vesicle by displaying the intersection of the Lagrangian Biomechanical Map and the eye vesicle. The cell contours are approximated by a sphere positioned on the detected nuclear center. Color code as in Supplementary Fig. 14. Embryo z1, anterior to the top, upper sections removed down to 64 μm below the embryo surface. (A) view from the outside. (B) View from the inside.

**Supplementary Table 1.** Image acquisition parameters. Recapitulates imaging conditions for the different specimens analyzed here. The ID column provides the identification specified in the BioEmergences database. Database repository link: http://bioemergences.eu/kinematics/

| Embryo | ID | Voxel | Voxel<br>Size<br>($\mu\text{m}^3$) | Timestep | Duration<br>(hpf) | T° (°C) | T° control |
| --- | --- | --- | --- | --- | --- | --- | --- |
| **z1** | 071226a | 512x512x120 | 1.37 | 2'33'' | 5.5-15 | 26 | 21° room |
| **z2** | 140523aF | 512x512x177 | 1.26 | 2'28'' | 5-13.8 | 26 | Okolab |
| **z3** | 141106aF | 512x512x148 | 1.38 | 2'24'' | 6-18 | 26 | Okolab |
| **z4** | 141108aF | 512x512x174 | 1.38 | 2'26'' | 6.5-20 | 28.6 | Okolab |
| **z5** | 141108a | 512x512x172 | 1.37 | 2'25'' | 6.5-20 | 24.7 | Okolab |

**Supplementary Table 2.** Tensor invariants computation. Left: name of descriptor computed as an invariant of a tensor. Right: mathematical definition of the descriptor. Given a tensor **A** expressed as a diagonal matrix with eigenvalues (*a*_1_, *a*_2_, *a*_3_). The first invariant is defined as the trace of the tensor: *I*_1_ = tr(**A**) = *a*_1_*+ a*_2_ *+ a*_3_ (as *P* and *MIC1*). The second invariant is defined as: 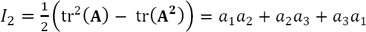 (as *Q*_*s*_, *Q*_*d*_ and *MIC2*). Third invariant is defined as the determinant: *I*_3_ = det(**A**) = *a*_1_*a*_2_*a*_3_ (as *J*). The rotation discriminant *D* is a descriptor that determines the limits in the presence of rotation (as described in *26*). This rotation descriptor has been preferred over the rotation rate provided by the skew-symmetric part **Ω** of the tensor **h** (*26*) or the second invariant *Q* of **h** used as vorticity descriptor in non-compressible fluids, as *D* provides a univocal metric of rotation in compressible flows *(P≠0)*. Interpretation of these invariants change according to the tensor and the temporal lengthscale of the deformation (Supplementary Table 3, S4).

|  |  |
| --- | --- |
| $P$ | $\text{tr}(\mathbf{h}) = h_1 + h_2 + h_3$ |
| $Q$ | $h_1 h_2 + h_2 h_3 + h_3 h_1$ |
| $D$ | $D^3 + PD^2 + QD + R = 0$ |
| $Q_s$ | $\varepsilon_1 \varepsilon_2 + \varepsilon_2 \varepsilon_3 + \varepsilon_3 \varepsilon_1$ |
| $Q_d$ | $d_1 d_2 + d_2 d_3 + d_3 d_1$ |
| $J$ | $\det(\mathbf{F})$ |
| $MIC1$ | $\text{tr}(\tilde{\mathbf{C}}) = \tilde{c}_1 + \tilde{c}_2 + \tilde{c}_3$ |
| $MIC2$ | $\tilde{c}_1 \tilde{c}_2 + \tilde{c}_2 \tilde{c}_3 + \tilde{c}_3 \tilde{c}_1$ |

**Supplementary Movie 1.**
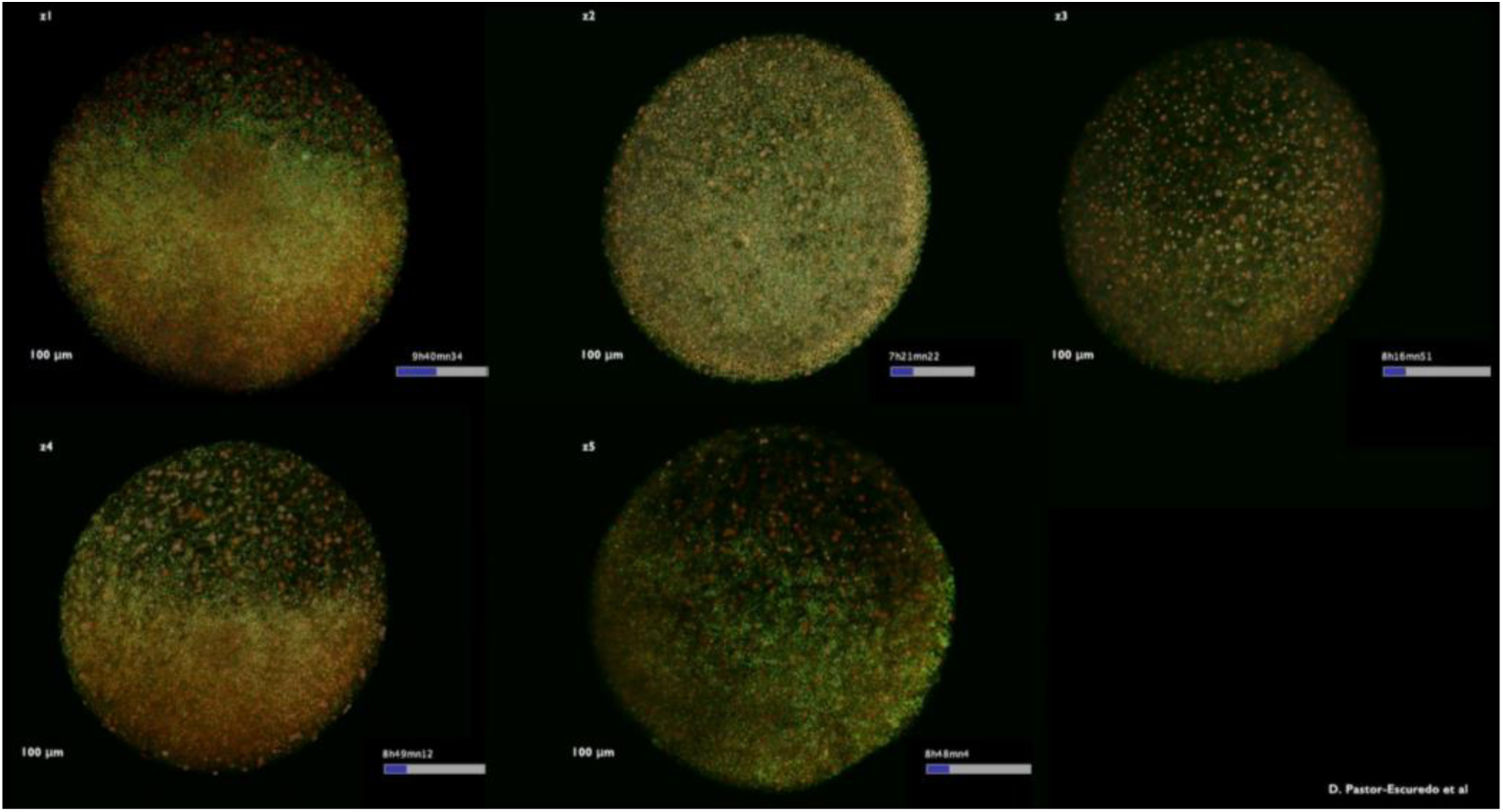
3D+time imaging of zebrafish gastrulation. Rendering of raw data overlaying the two channels for cell membranes (green) and nuclei (red) respectively (Mov-IT software). Embryos z1 to z5 indicated top left. Nuclei are marked with H2B-mCherry and membranes are marked with eGFP-HRAS. Embryos are aligned temporally to roughly synchronize the onset of their development. They are imaged from the animal pole and observed first from the outside then from the inside. High-resolution movie available at: http://bioemergences.eu/kinematics/movies/MovieS1.mp4

**Supplementary Movie 2.**
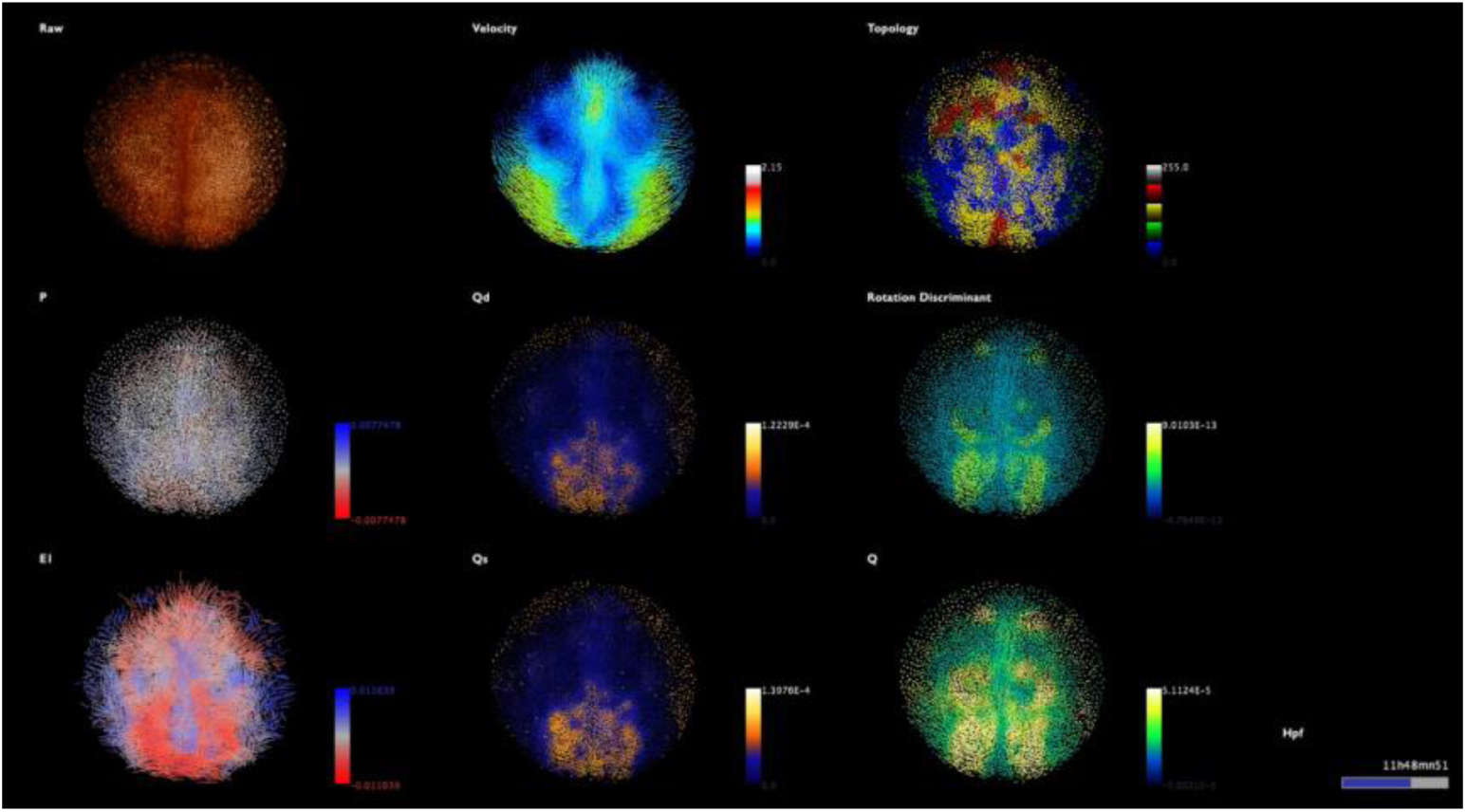
Eulerian descriptors (embryo z1) Instantaneous kinematics description from mid-gastrulation to early segmentation (8-14 hpf), embryo z1 imaged from the animal pole, outside view then inside view. Raw data top left, descriptor indicated top left of each panel. Color code as in Fig. 1. Scale bar 100 μm. Temporal scale bottom right. High-resolution movie available at: http://bioemergences.eu/kinematics/movies/MovieS2.mp4

**Supplementary Movie 3.**
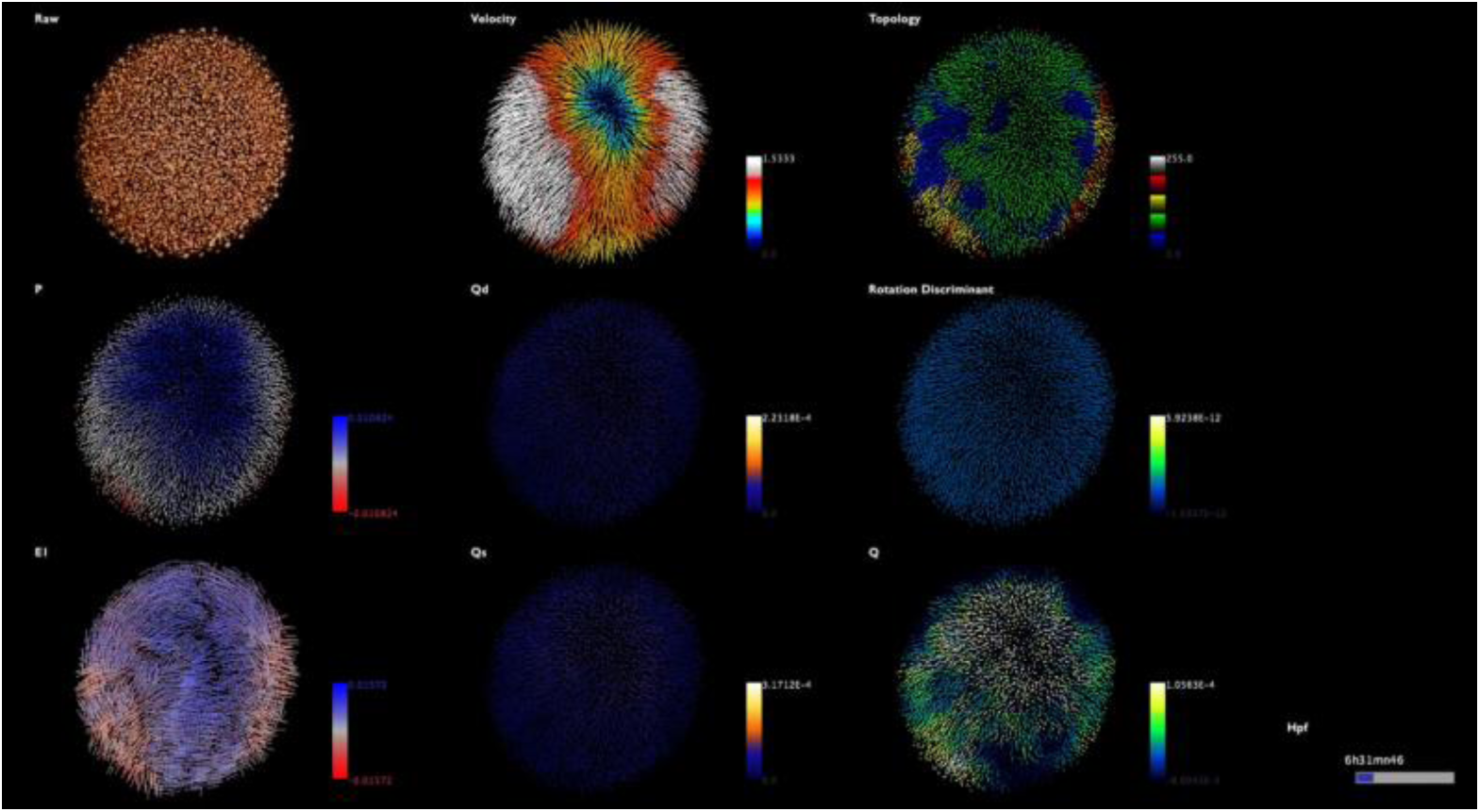
Eulerian descriptors (embryo z2) Instantaneous kinematics description from mid-gastrulation to early segmentation (8-14 hpf) embryo z2 imaged from the animal pole, outside view then inside view. Raw data top left, descriptor indicated top left of each panel. Color code as in Fig. 1. Scale bar 100μm. Temporal scale bottom right. High-resolution movie available at: http://bioemergences.eu/kinematics/movies/MovieS3.mp4

**Supplementary Movie 4.**
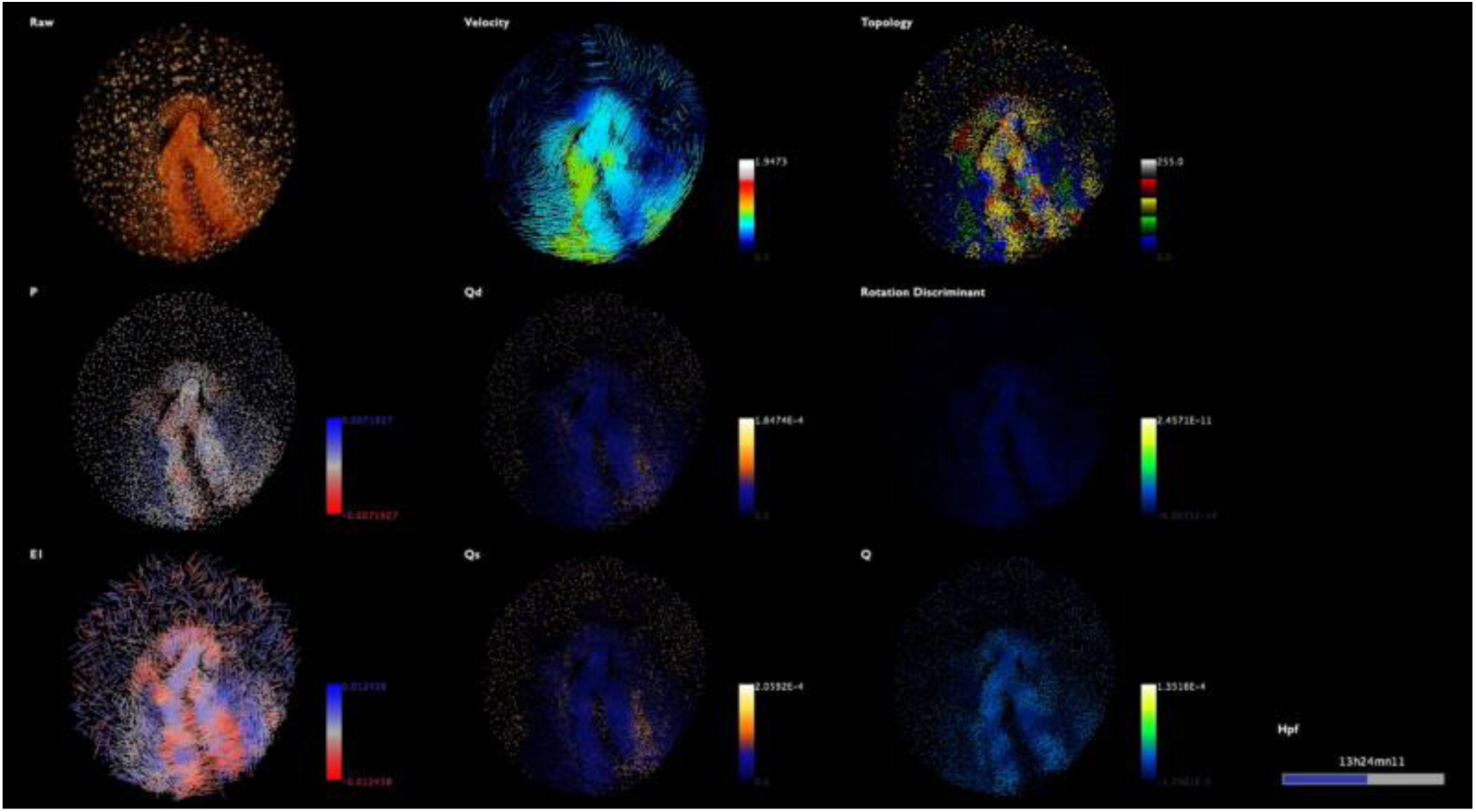
Eulerian descriptors (embryo z3) Instantaneous kinematics description from mid-gastrulation to early segmentation (8-14 hpf) embryo z3 imaged from the animal pole, outside view then inside view. Raw data top left, descriptor indicated top left of each panel. Color code as in Fig.1. Scale bar 100μm. Temporal scale bottom right. High-resolution movie available at: http://bioemergences.eu/kinematics/movies/MovieS4.mp4

**Supplementary Movie 5.**
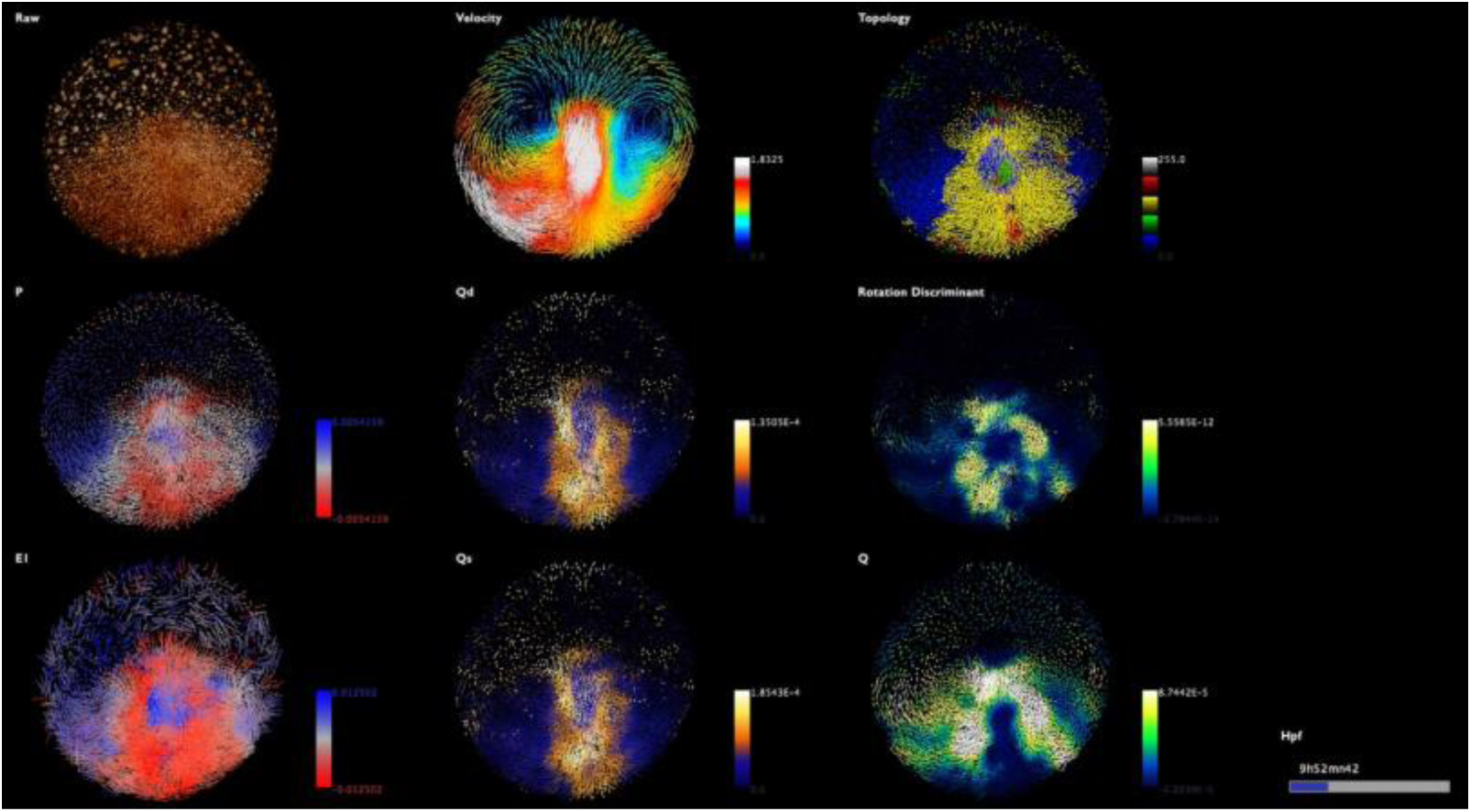
Eulerian descriptors (embryo z4) Instantaneous kinematics description from mid-gastrulation to early segmentation (8-14 hpf) embryo z4 imaged from the animal pole, outside view then inside view. Raw data top left, descriptor indicated top left of each panel. Color code as in Fig.1. Scale bar 100μm. Temporal scale bottom right. High-resolution movie available at: http://bioemergences.eu/kinematics/movies/MovieS5.mp4

**Supplementary Movie 6.**
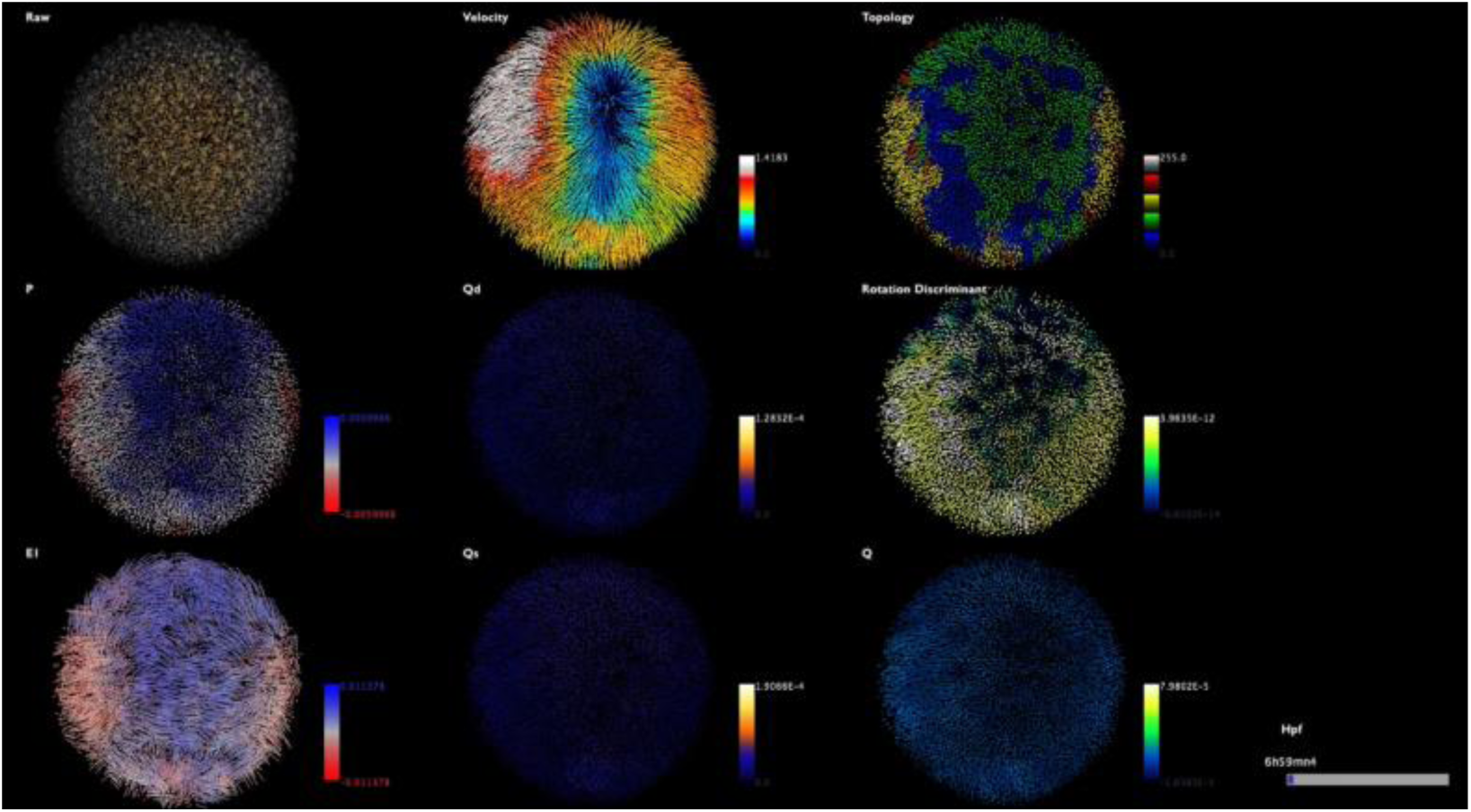
Eulerian descriptors (embryo z5) Instantaneous kinematics description from mid-gastrulation to early segmentation (8-14 hpf) embryo z5 imaged from the animal pole, outside view then inside view. Raw data top left, descriptor indicated top left of each panel. Color code as in Fig. 1. Scale bar 100 μm. Temporal scale bottom right. High-resolution movie available at: http://bioemergences.eu/kinematics/movies/MovieS6.mp4

**Supplementary Movie 7.**
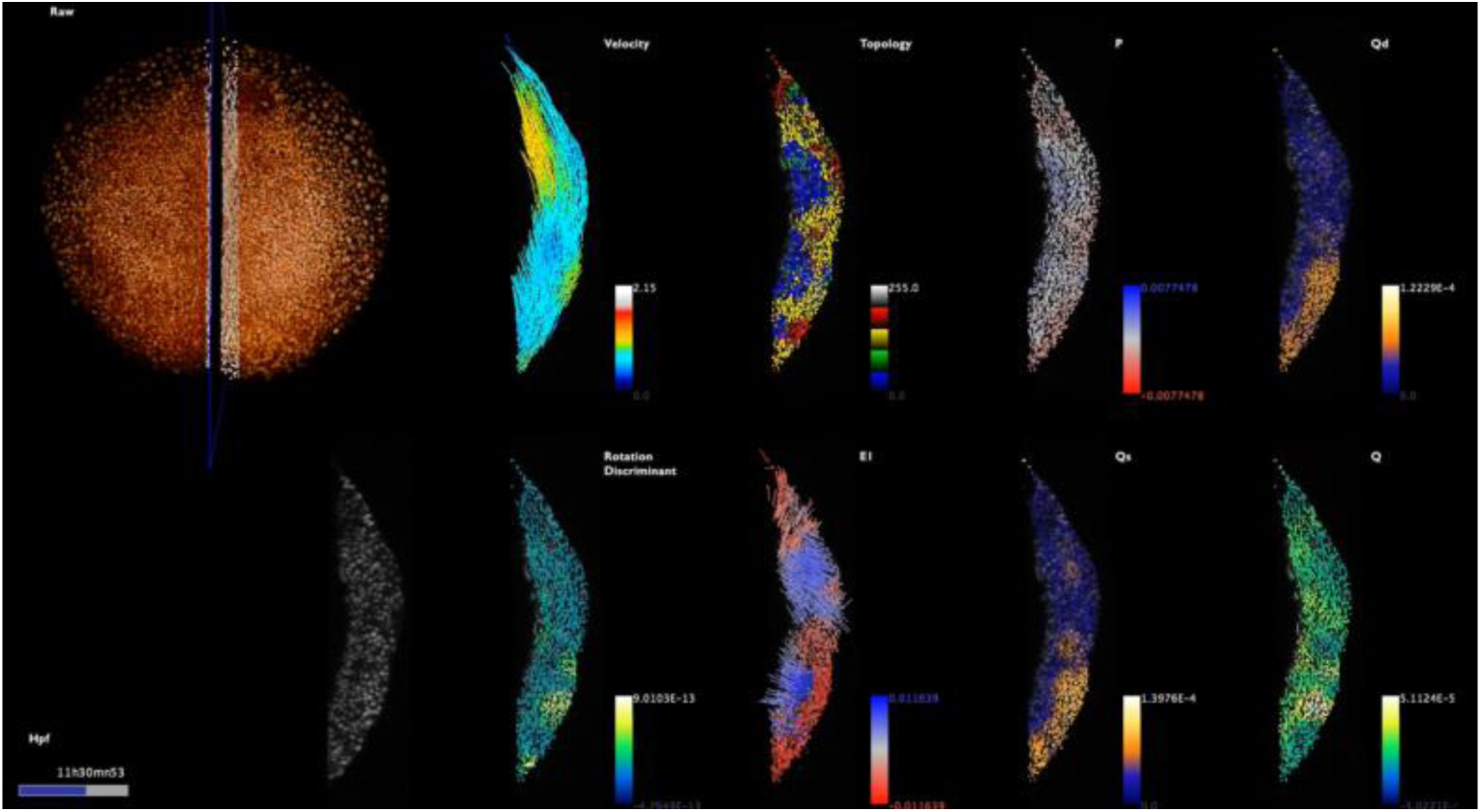
Deformation rates at the midline (embryo z1) Instantaneous kinematics description from mid-gastrulation to early segmentation (8-14 hpf) embryo z1 imaged from the animal pole. Raw data top left, outside view then inside view. The midline is marked by an orthoslice (in black) and about 3-5 row of cells are selected (white) on each side of the midline for a lateral visualization (bottom left). Descriptors as in Supplementary Fig. 4. indicated top right of each panel. Color code as in Fig. 1. Scale bar 100 μm. Temporal scale bottom right. High-resolution movie available at: http://bioemergences.eu/kinematics/movies/MovieS7.mp4

**Supplementary Movie 8.**
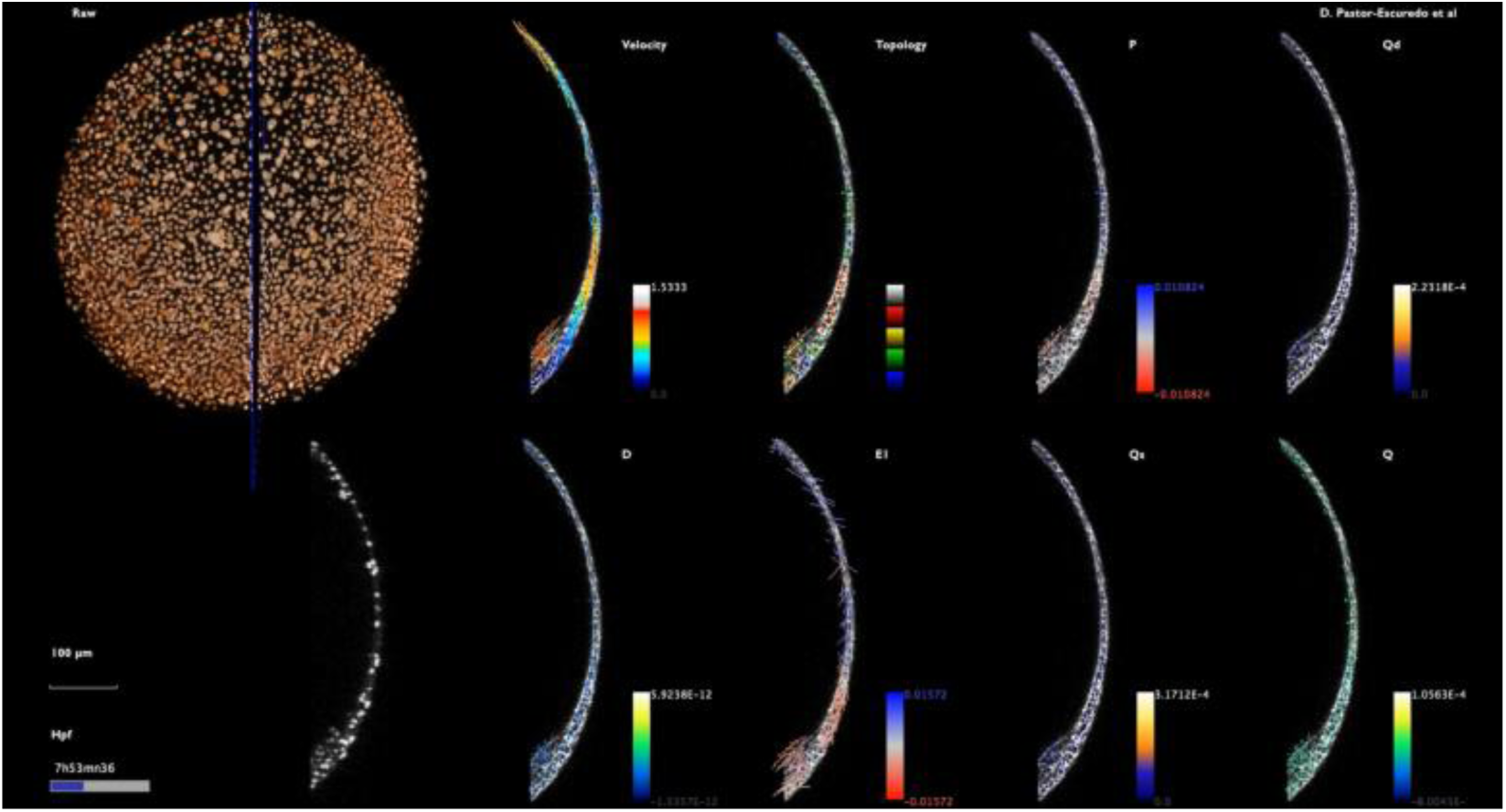
Deformation rates at the midline (embryo z2) Same as Supplementary Movie 7 for embryo z2. High-resolution movie available at: http://bioemergences.eu/kinematics/movies/MovieS8.mp4

**Supplementary Movie 9.**
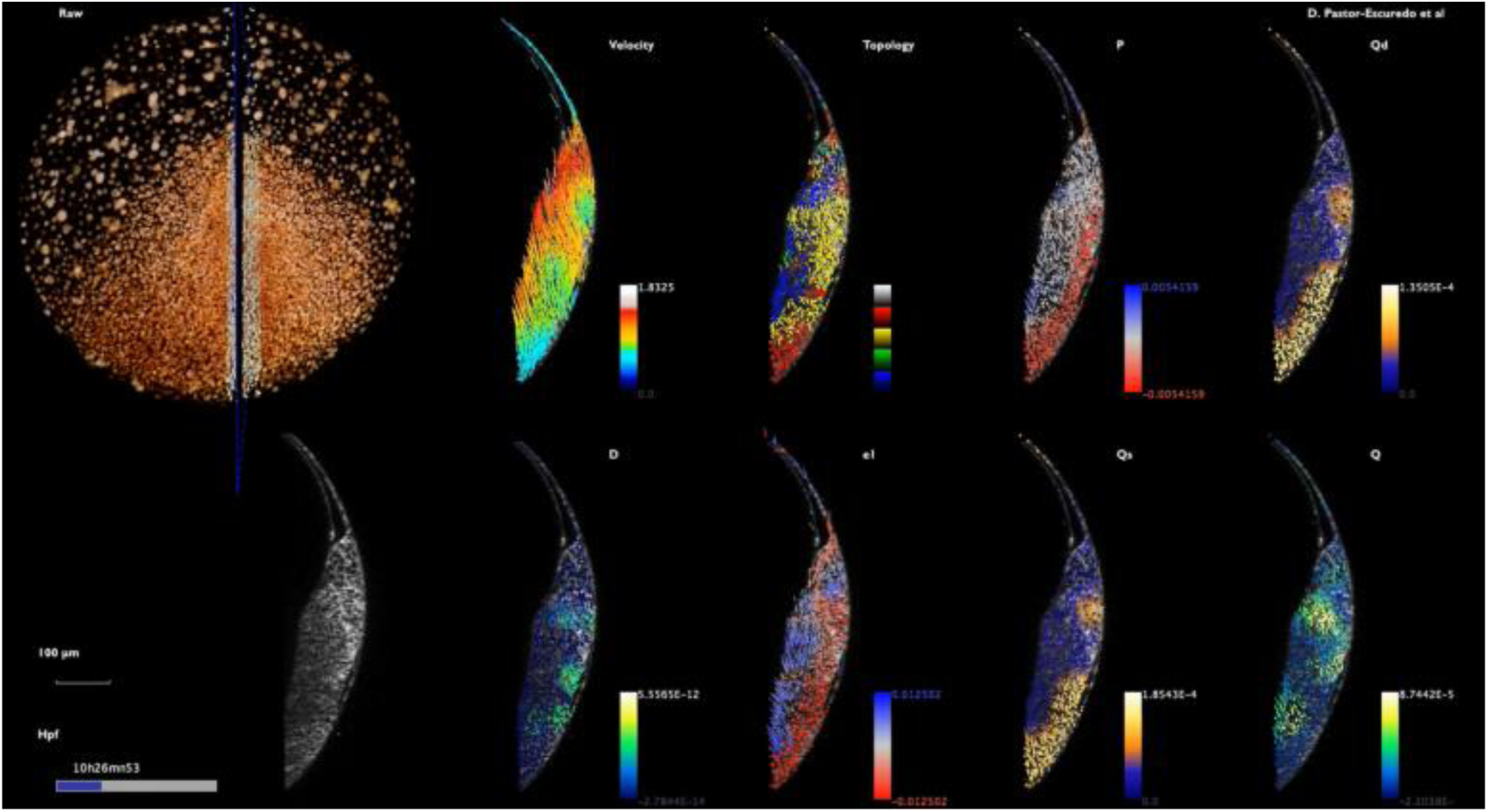
Deformation rates at the midline (embryo z3) Same as Supplementary Movie 7 for embryo z3. High-resolution movie available at: http://bioemergences.eu/kinematics/movies/MovieS9.mp4

**Supplementary Movie 10.**
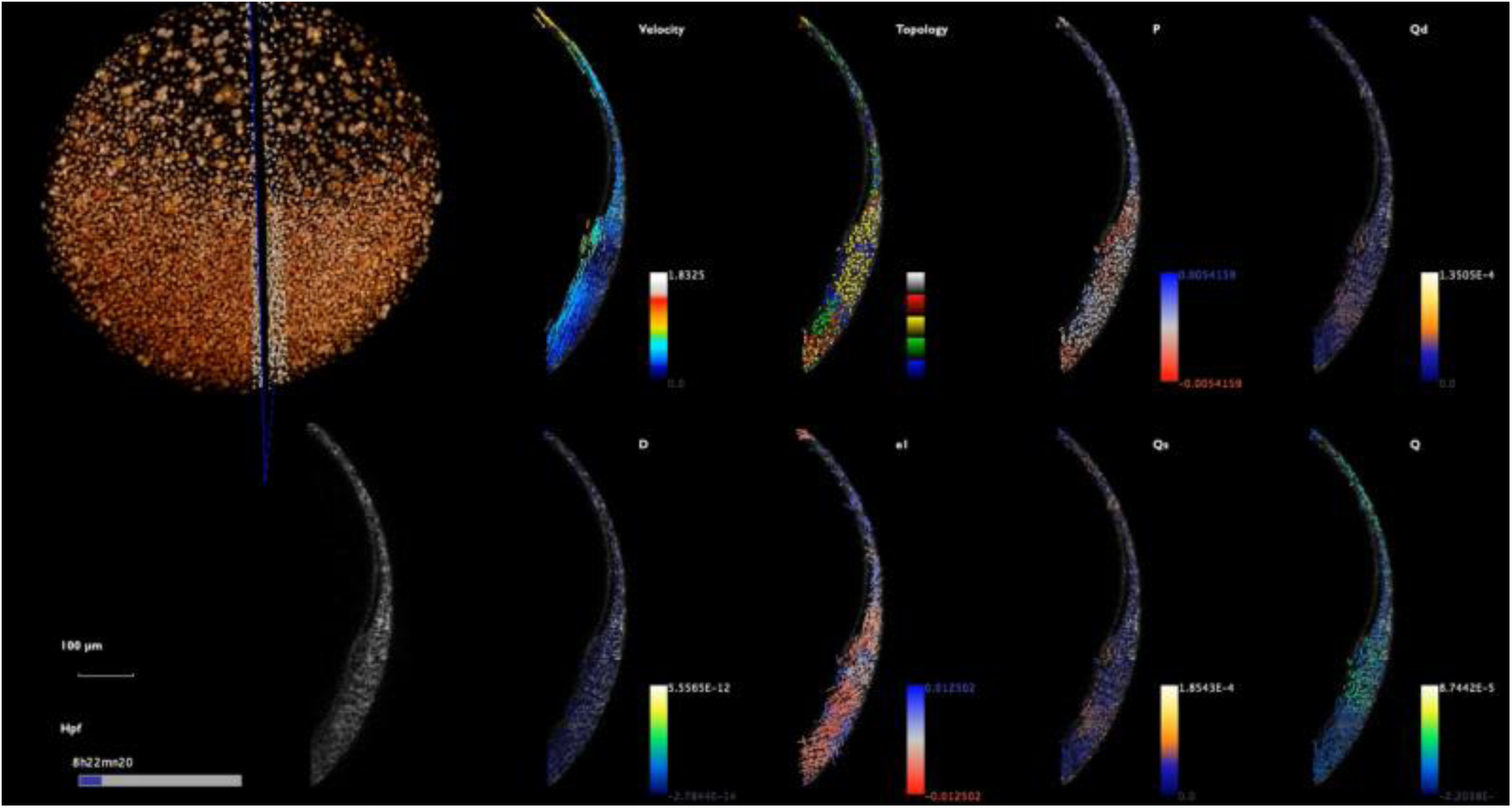
Deformation rates at the midline (embryo z4) Same as Supplementary Movie 7 for embryo z4. High-resolution movie available at: http://bioemergences.eu/kinematics/movies/MovieS10.mp4

**Supplementary Movie 11.**
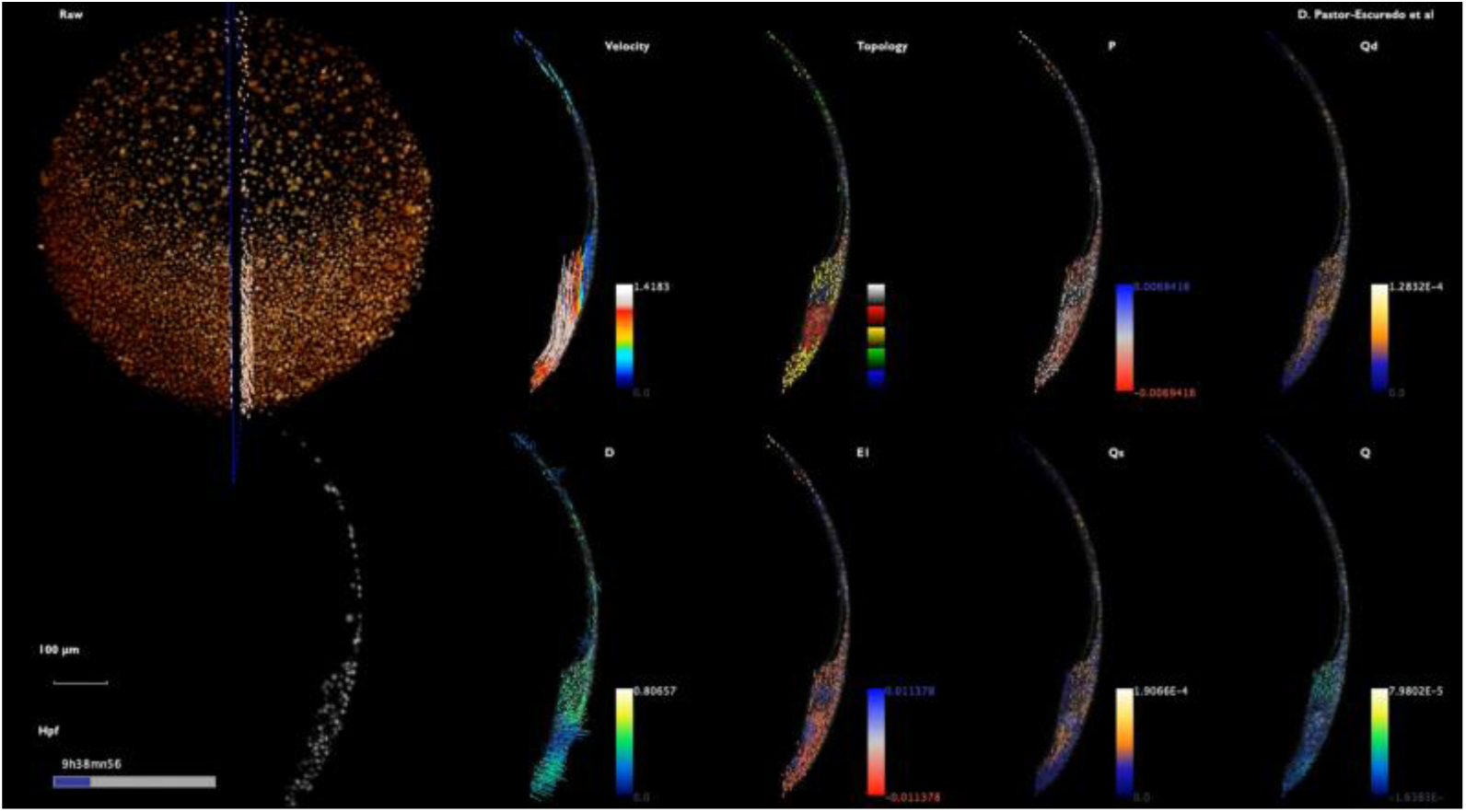
Deformation rates at the midline (embryo z5) Same as Supplementary Movie 7 for embryo z5. High-resolution movie available at: http://bioemergences.eu/kinematics/movies/MovieS11.mp4

**Supplementary Movie 12.**
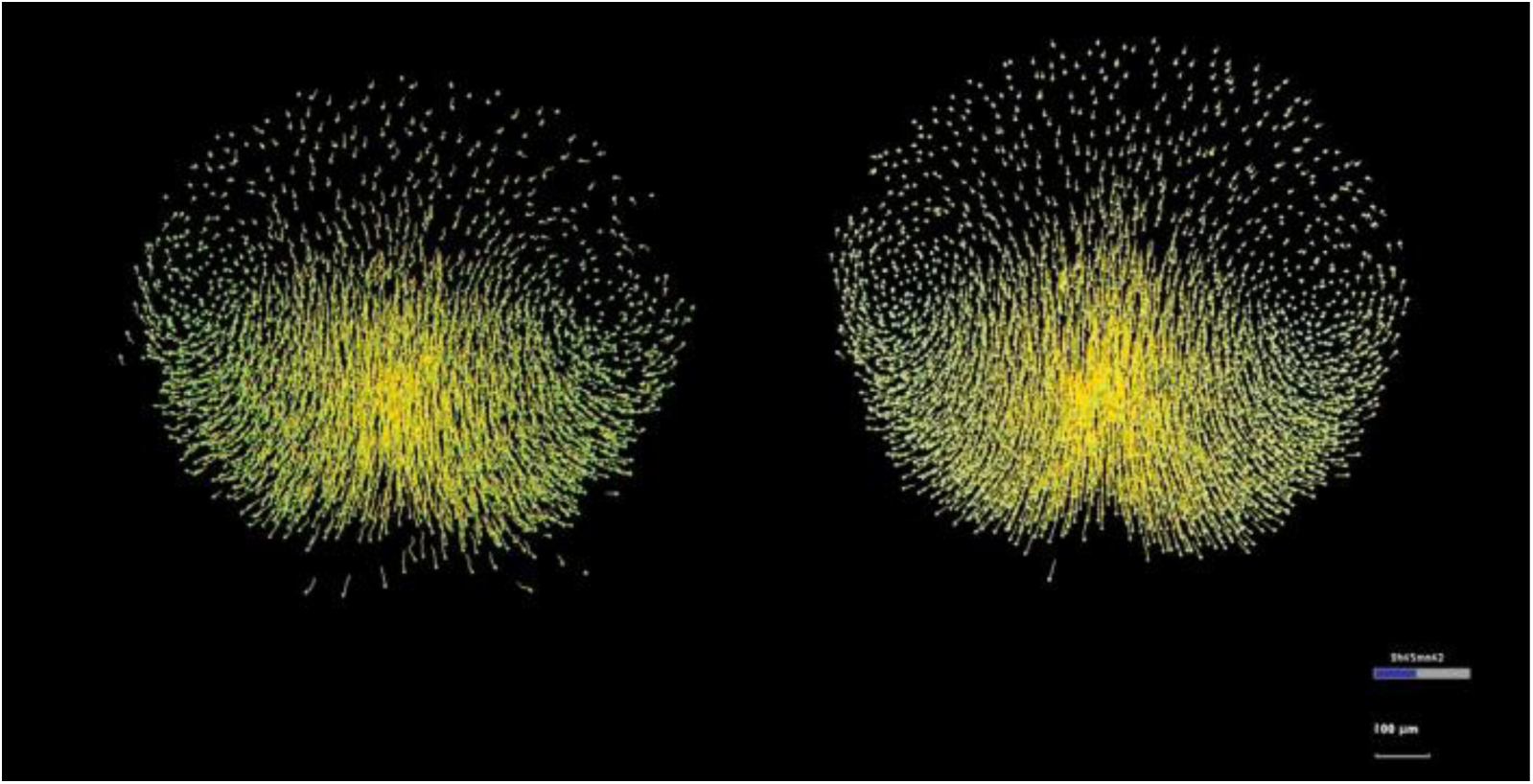
Flow approximation of the cell lineage. Embryo z1 animal pole view (Mov-IT software). Displacements implicit in the digital cell lineage (left) compared to displacements in the flow field (right) after the temporal smoothing and the spatial regularization of the cell displacements. In the left panel, the cells that get into the imaged volume are excluded. The flow field is composed of complete trajectories resulting from the integration of the displacements. It produces an artifact, keeping in the field of view cells that normally moved out of it. It should be noted that this artifact is solved by working with a selection of cells that are kept in the imaged volume for the whole imaging sequence (shield selection as in Supplementary Fig. 8). High-resolution movie available at: http://bioemergences.eu/kinematics/movies/MovieS12.mp4

**Supplementary Movie 13.**
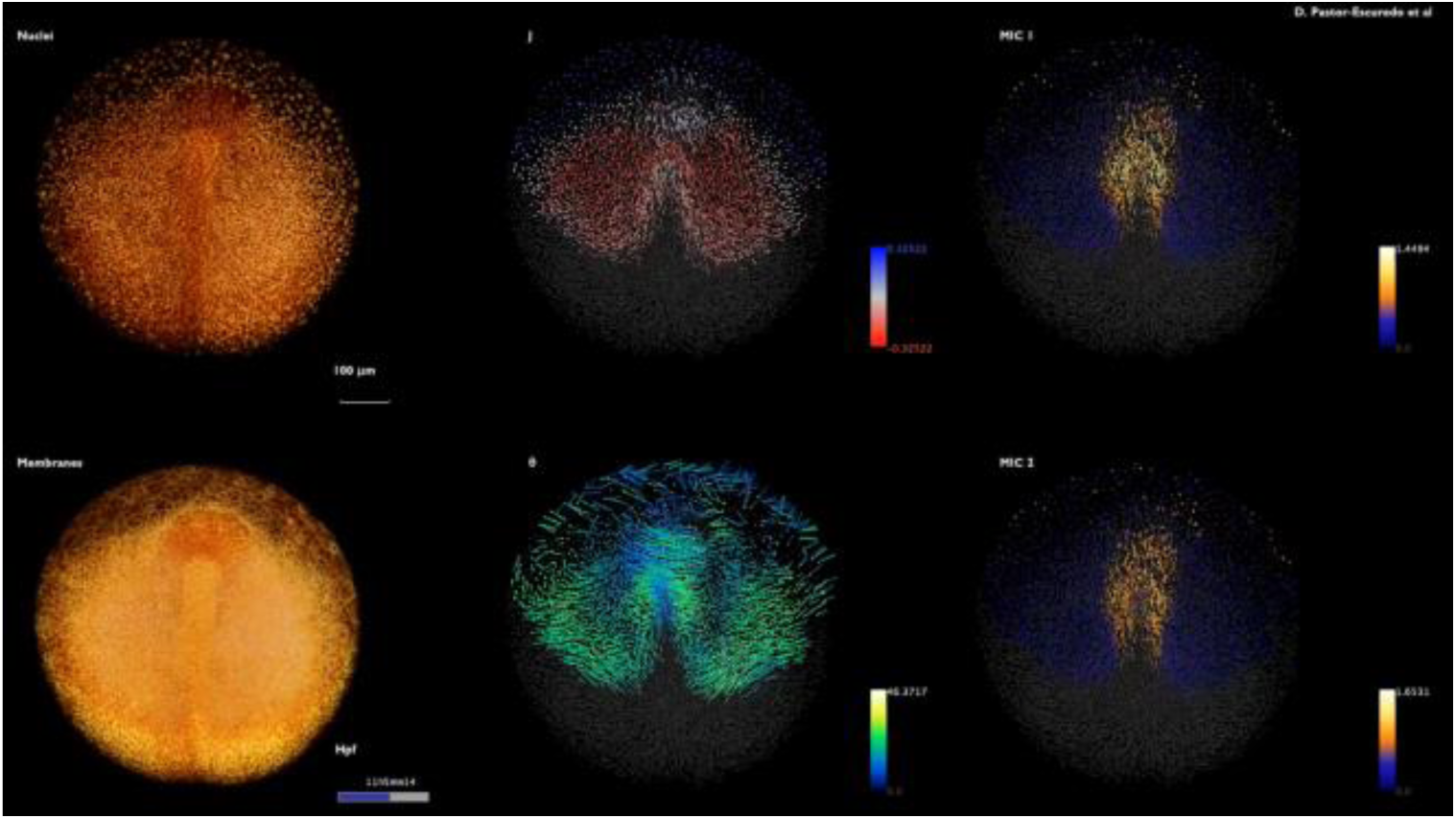
Maps of cumulative LBPs. Cumulative LBPs displayed as maps from mid-gastrulation to early segmentation (8-14 hpf) embryo z1 imaged from the animal pole. The cumulative LBPs are computed with *t*_ini_=8 hpf. First view from the inside then outside then inside. Raw data nuclei top left, membranes bottom left. Descriptor indicated top left of each panel. Color code as in Fig. 2. Scale bar 100 μm. Temporal scale bottom right of the membrane raw data panel. As in Supplementary Movie 12, new cells that get into the field of view are not seen. High-resolution movie available at: http://bioemergences.eu/kinematics/movies/MovieS13.mp4

**Supplementary Movie 14.**
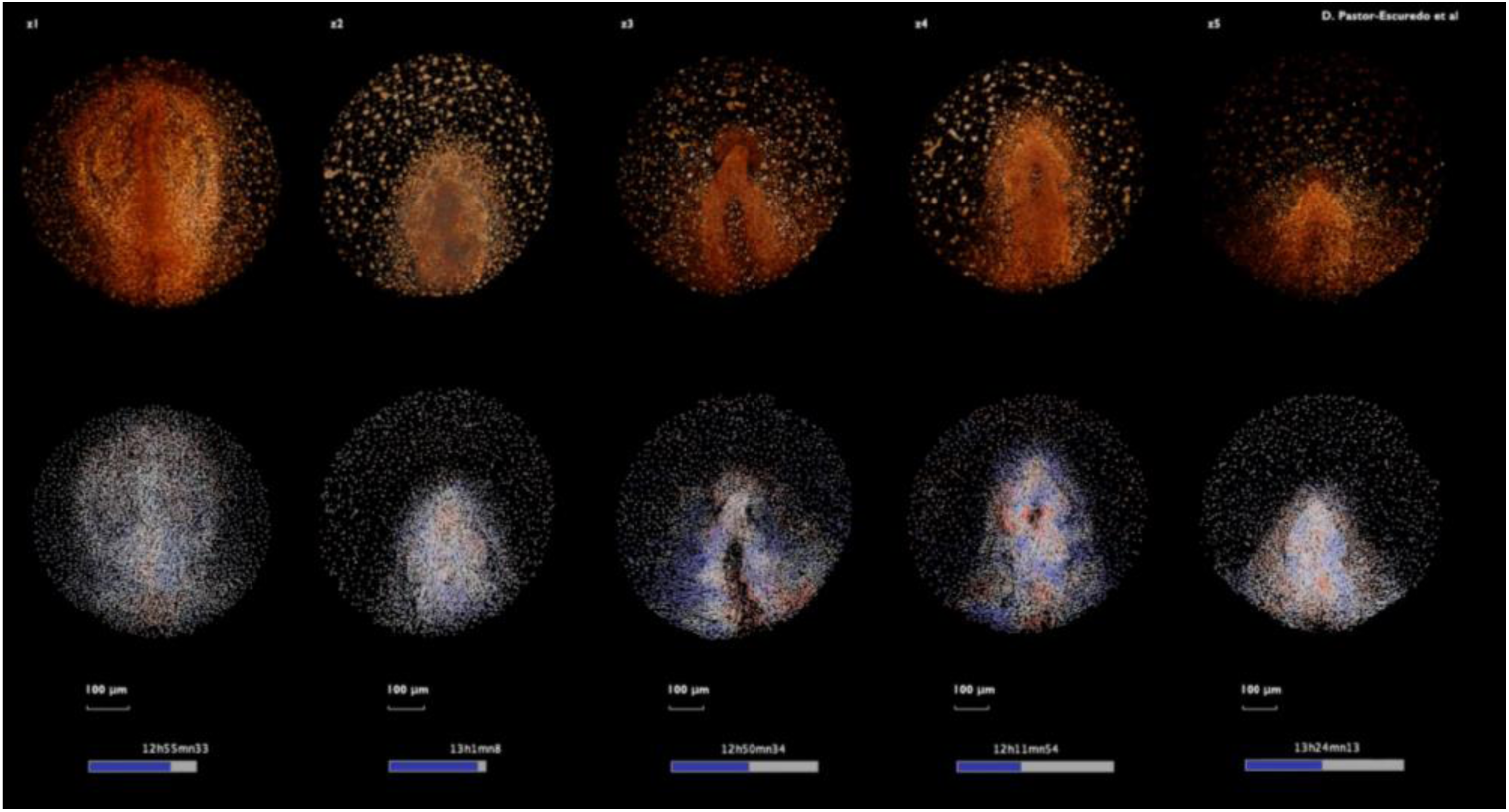
Comparison of instantaneous compression and expansion. First row: 3D rendering of raw nuclei for z1-z5 (indicated top left of each column). Second row: descriptor *P* with color code as in Fig. 1. Embryos have been positioned as in Supplementary Movie 1 and temporally aligned according to Fig. 3. View first from inside, then outside and inside. Scale bar 100 μm. Temporal scale bottom of each column. High-resolution movie available at: http://bioemergences.eu/kinematics/movies/MovieS14.mp4

**Supplementary Movie 15.**
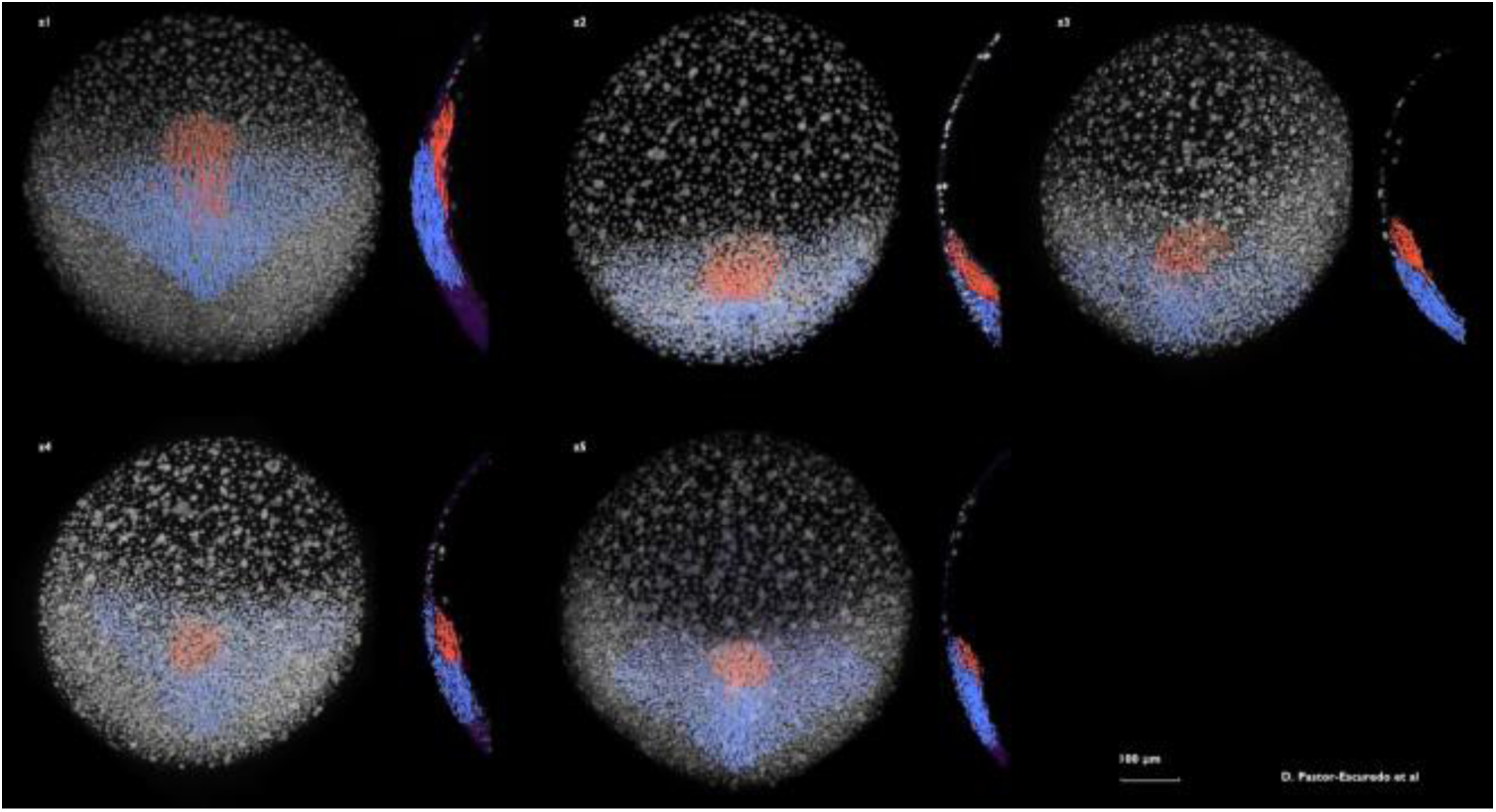
Tailbud selection of germ layers. Render of germ layers: epiblast (blue) and hypoblast (red). Selections of both populations have been done by the end of gastrulation for each dataset (tailbud selection as in Supplementary Fig. 7) and propagated through the trajectories along the whole interval of analysis (7-14 hpf). As the field of view and orientation are not exactly the same, the hypoblast appears at different time points in each movie. From left-right, up-down: z1, z2, z3, z4 and z5. High-resolution movie available at: http://bioemergences.eu/kinematics/movies/MovieS15.mp4

**Supplementary Movie 16.**
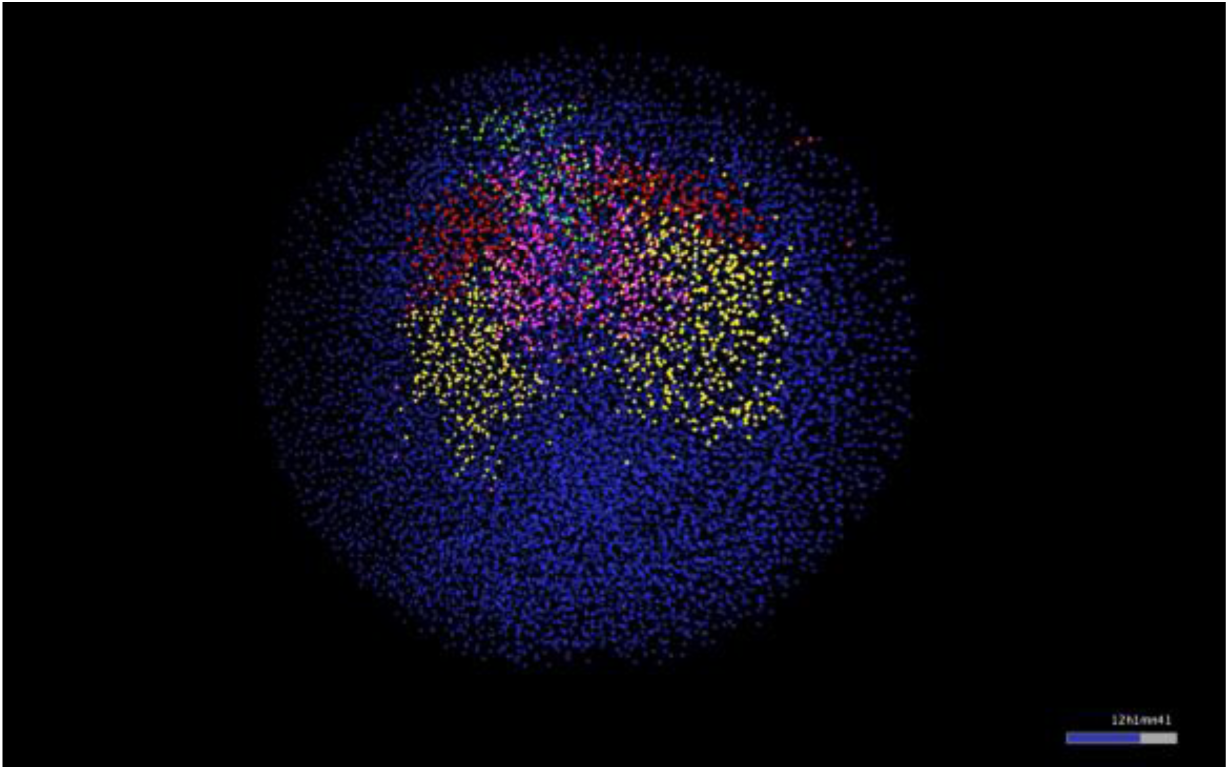
Lagrangian Biomechanical Map. Spatio-temporal distribution of the mechanical domains (as in Fig. 4) from 8 to 13 hpf. Outside view of embryo z1, animal pole, anterior to the top. The four biomechanical domains are obtained by clustering the selected cell population (shield selection as in Supplementary Fig. 8) at 8 hpf into four domains according to the mechanical signature of the trajectories. Cells are labeled by 8 hpf according to their respective biomechanical domain and labels are propagated along the cell lineage. Visualization with Mov-IT software. High-resolution movie available at: http://bioemergences.eu/kinematics/movies/MovieS16.mp4

**Supplementary Movie 17.**
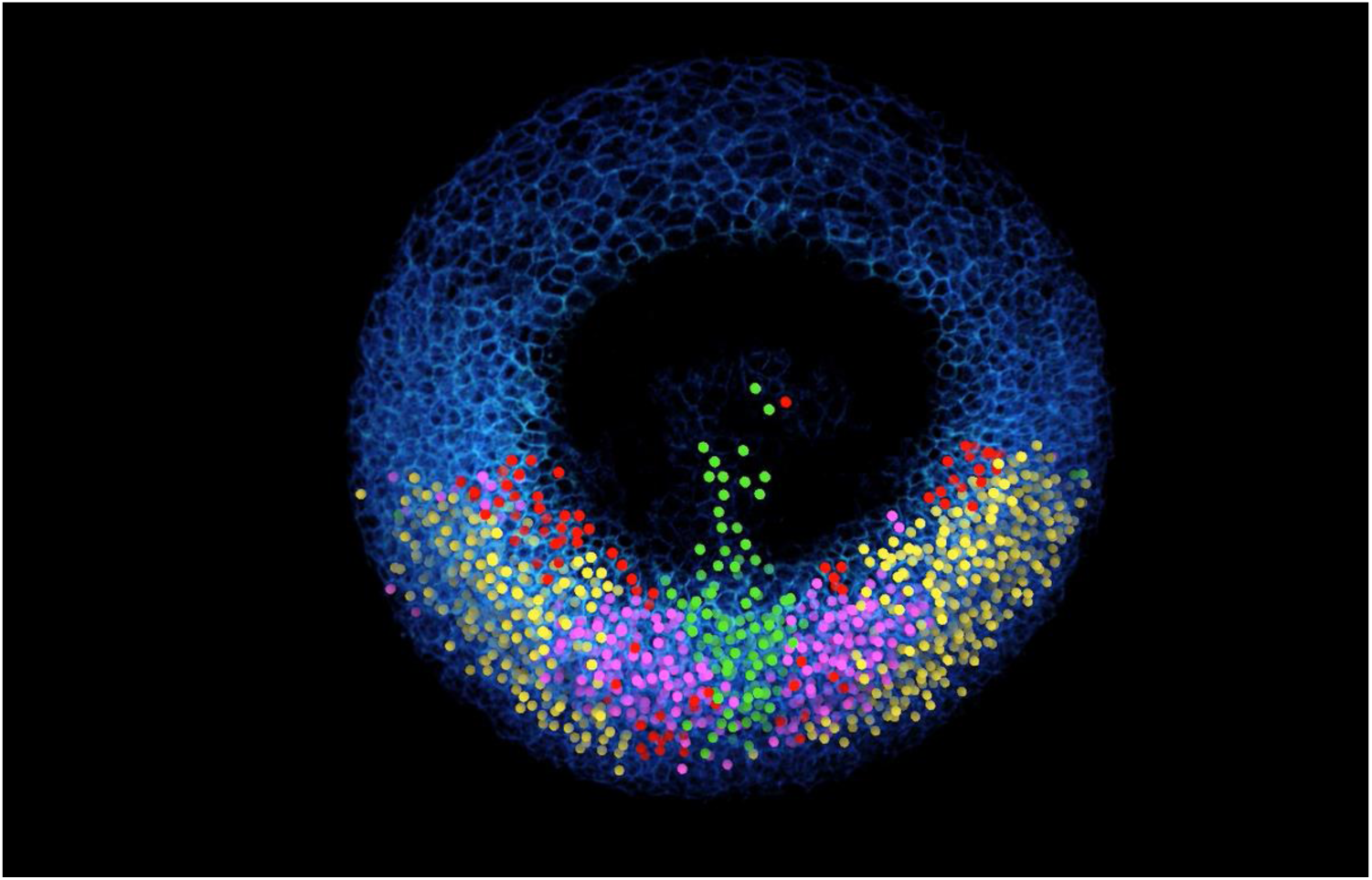
Lagrangian Biomechanical Map section. Same as S16 but removing upper section down to 65 μm below the embryo surface. High-resolution movie available at: http://bioemergences.eu/kinematics/movies/MovieS17.mp4

**Supplementary Movie 18.**
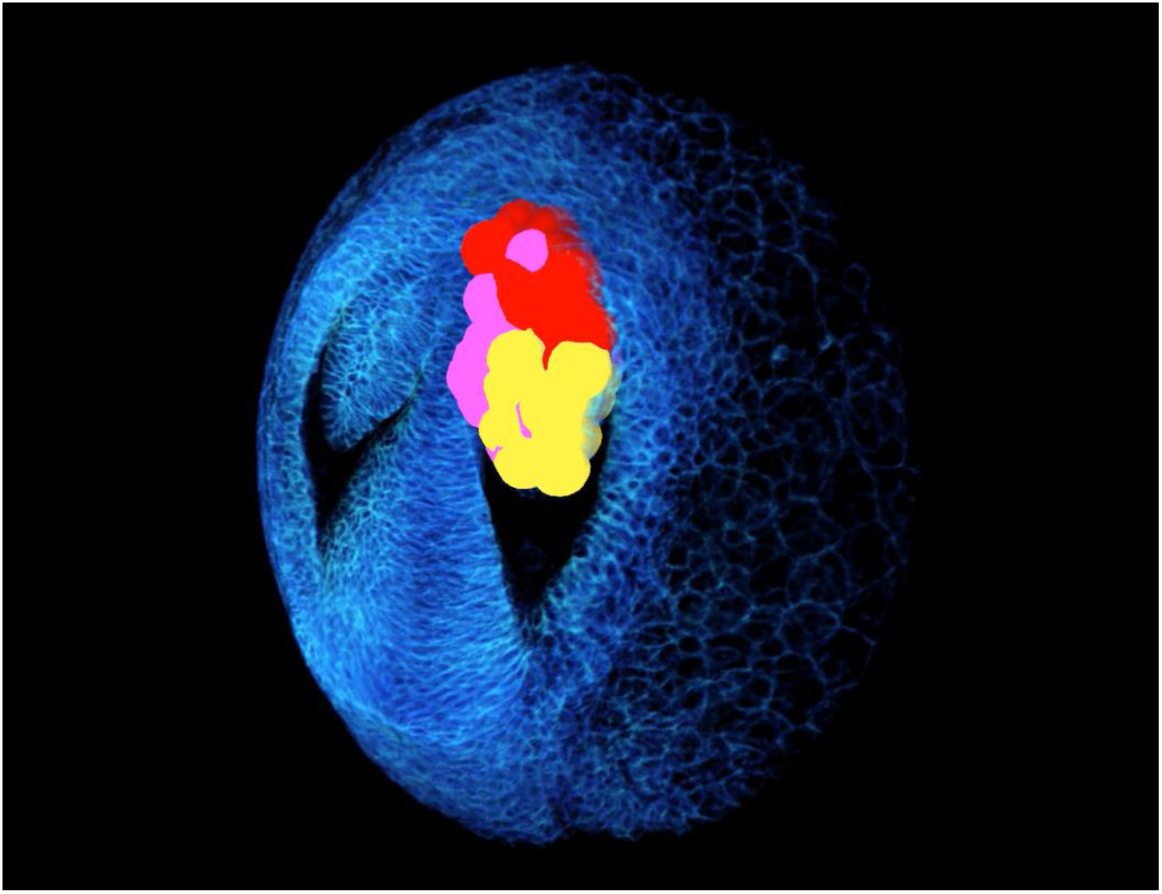
Lagrangian Biomechanical Map of the eye vesicle. As in Supplementary Fig. 14. Highlights the biomechanical domains in the eye vesicle by displaying the intersection of the Lagrangian Biomechanical Map and the eye vesicle. A sphere positioned at the detected nuclear center approximates the cell contours. Color code as in Supplementary Fig. 13. Embryo z1, anterior to the top, upper sections sequentially removed down to 65 μm below the embryo surface. High-resolution movie available at: http://bioemergences.eu/kinematics/movies/MovieS18.mp4

